# Phenotypic and transcriptomic similarity between the N2 Ancestral and a tropical wild isolate of *C. elegans* reveals divergence from the reference Bristol strain

**DOI:** 10.1101/2025.06.27.662035

**Authors:** Jaqueline Hersch-González, Nalley Cano-Domínguez, Mariana Zurita-León, Maribel Soto-Nava, Santiago Avila-Rios, Victor Julian Valdes

## Abstract

In recent years, the scientific community has increasingly recognized the importance of incorporating ecologically relevant perspectives into laboratory research. In the case of the free-living nematode *Caenorhabditis elegans*, numerous studies have documented the domestication of the N2 Bristol strain (isolated in 1951). This has led to a growing interest in recently isolated wild strains from diverse latitudes, which offer insights into natural variation evolution and life-history traits. Here, we compared a recently isolated tropical strain from Mexico City to the N2 Bristol strain. To contextualize laboratory adaptation, we also included the N2 Ancestral strain, a cryopreserved lineage from 1969 with minimal generational drift. Phenotypic assays revealed that, under standard laboratory conditions, the Mexican strain exhibited reduced lifespan and fertility, but enhanced resistance to *Pseudomonas aeruginosa*, whereas the Ancestral strain showed higher oxidative stress tolerance but reduced thermotolerance. RNA-seq analyses showed that transcriptomic profiles of the Mexican and Ancestral strains were more similar to each other than to the N2 Bristol, suggesting that long-term domestication has driven regulatory divergence. Differential gene expression analyses identified strain-specific signatures in stress, immune and collagen-related pathways. Under heat stress, transcriptional profiling revealed that only a small set of canonical heat shock genes was commonly upregulated across the three strains, yet wild strains showed more dynamic regulation, while N2 Bristol exhibited a distinct, possible preconditioned response. These findings reveal phenotypic trade-offs and regulatory divergence shaped by natural versus laboratory environments, and underscore evolutionary dynamics and adaptive potential of *C. elegans* in response to distinct ecological histories.

## Introduction

The free-living nematode *Caenorhabditis elegans* has been an extraordinary model organism for over 70 years advancing research in molecular biology, development, aging, neuroscience and evolution. It played a fundamental role in the discovery of apoptosis, microRNAs and RNA interference, as was the first multicellular organism to have its entire genome sequenced (Ellis and Horvitz 1986; White el al. 1986; Kenyon et al. 1993; Wightman et al. 1993; Lee et al. 1993; Fire et al. 1998; C. elegans Sequencing Consortium 1998). *C. elegans* was first identified in Algeria in 1900 by Emile Maupas and later studied in the 1940s by Victor Nigon and Ellsworth Dougherty (Nigon and Félix 2017). However, the widespread adoption of *C. elegans* as a model organism was largely driven by the extraordinary vision of Sydney Brenner, who in the 1960s started working with the genetics of a *C. elegans*’s strain isolated in 1951 in Bristol, UK, that he later designated as the “N2 Bristol” wild-type (WT) strain (Brenner 1974; Sterken et al. 2015). Today, hundreds of laboratories worldwide use *C. elegans* as a model system, with the N2 Bristol strain serving as a reference that has been extensively characterized and genetically manipulated (Ankeny, 2001). Remarkably, a tube presumed to contain the original N2 Bristol strain frozen in 1969 was thawed in 1980 to establish what is now considered the “Ancestral” N2 stock, available through the *Caenorhabditis* Genetics Center (CGC). It is estimated that this ancestral strain is less than six generations away from the one used by Brener and frozen in 1969 (Sterken et al. 2015).

Part of the success of *C. elegans* as a model organism is based on its easy cultivation and maintenance: it can be grown on agar plates at 20 °C (±5 °C) with a lawn of *E. coli* as a food source (Corsi et al. 2015). These standardized laboratory conditions have been implemented by most laboratories worldwide. However, they differ significantly from the complex and dynamic natural environments that *C. elegans* encounters in the wild: fluctuation in temperature, oxygen and humidity, along with scarcity yet diversity of food sources within a tridimensional habitat where it faces pathogens and predators. As a result, wild nematodes must display substantial phenotypic and transcriptional flexibility to survive.

Although impossible to calculate, given the short-life cycle of *C. elegans* (one generation every 72h at 20 °C), it is reasonable to assume that thousands of generations of the N2 Bristol strain have been propagated under laboratory conditions since its initial isolation 70 years ago. Consequently, it is also possible to assume that a process of adaptation and domestication to laboratory conditions, along with unintended genetic drift, have occurred over this period in the N2 Bristol strain ( Weber et al. 2010; Sterken et al. 2015). In recent years, efforts have been made to isolate and study different wild isolates from diverse geographical locations. For instance, the *C. elegans* Natural Diversity Resource (CaeNDR) currently catalogs over 1700 wild isolates collected from every continent except Antarctica (Crombie et al. 2024). Many of these recently isolated strains exhibit both genotypic and phenotypic differences compared to each other and to the standar N2 Bristol strain (Weber et al. 2010; Sterken et al. 2015; Zhao et al. 2018; Urban et al. 2021). One illustrative example of natural variation in behavior is the “social” phenotype observed in a Hawaiian isolate and other wild strains, which is associated with a different isoform at the NPR-1 neuropeptide Y receptor. These strains carry a phenylalanine at position 215, whereas the standard N2 Bristol strain has a valine at this position. This variant leads to a preference for aggregation of individuals under high-oxygen laboratory conditions, in contrast to the “*solo*” feeding behavior displayed by the N2 Bristol strain on the agar plates (de Bono and Bargmann 1998; Gray et al. 2004; Sterken et al. 2015).

Understanding the phenotypic divergence between laboratory-adapted strains like N2 and wild isolates offers insights into the consequences of artificial selection and genetic drift in the experimental systems. Thus, in this study, we compared life-traits and gene expression profiles among three *C. elegans* strains: (i) a N2 Bristol strain recently acquired from the CGC (N2 lab), (ii) the ancestral N2 strain, also from the CGC, and (iii) a wild tropical strain recently isolated in Mexico City. We found that the strains respond differently to biotic and abiotic stressors, exhibiting differences in lifespan and fertility, and that N2 Lab strain display the most distinct transcriptomic profile, while the ancestral and the Mexican strains are more similar to each other. These findings underscore the importance of environmental context in shaping adaptation, and highlights the importance of considering strain history and adaptation when interpreting results, as these factors can significantly influence experimental outcomes and their broader implications.

## Results

### Isolation and characterization of a tropical wild *Caenorhabditis elegans* strain

While Bristol, UK, is a coastal city with a humid cold climate (2-25 °C; average 11°C), Mexico city is located over 2,000 meters above sea level characterized by well defined dry and wet seasons and a tropical temperature (8-34 °C; average 20°). To compare the N2 bristol strain with a tropical *C. elegans* isolate, we collected samples from rotting fruit in gardens across Mexico City, obtaining selfing worm populations many of which were compatible to *Oscheius*. Microscope observation revealed some individuals with the distinctive double-bulb pharynx characteristic of *C. elegans* (Figure 1A) that was clonally propagated. We confirmed the species identity by sequencing a 777 bp amplicon from the ribosomal ITS2 region by PCR (see methods, Barrière and Félix 2014) revealing 99.56% homology with a *C. elegans* strain genome (supplementary Abi file 1 and 2; Figure S1 A and B). We backcrossed this new Mexican isolate with males from the *C. elegans* OH10689 strain (carrying a neuronal RFP reporter), obtaining fluorescent, self-propagating progeny (data not shown). The Mexican strain was cultivated under standard laboratory conditions, but to preserve its characteristics as close to the original, several vials were frozen. For all the experiments in this study, we only employed worms within the first 10 generations derived from these original frozen stocks. The Mexican isolated strain was registered as VJV001 in the *Caenorhabditis* Genetics Center (CGC) and sent for genome sequencing at CaeNDR. In the rest of the text, it will be referred to as “Mexican strain”. Initial observations revealed that the Mexican strain deployed the “social feeding behavior”, meaning that worms remain clumped at the edges of the bacterial lawn (Figure S1 C), a region reported to have a lower oxygen concentration (Gray et al. 2004). Several wild *C. elegans* isolates share this phenotype which has been attributed to a different isoform of the NPR-1 neuropeptide Y receptor (215F) (de Bono et al. 1998). The presence of the “social” allele in the Mexican strain was corroborated by sequencing (data not shown). In parallel, we obtained a sample of the CGC’s N2 ‘ancestral’ strain that is estimated to be within six generations from the original Bristol first frozen stock, and it will be compared to the standard CGC’s N2 (var Bristol) strain obtained in 2023, here referred as “N2 Lab”. These three strains will allow us to assess the differences among a recent tropical isolate, the ancestral wild-type and the laboratory domesticated N2 Bristol strain. No differences were observed between the three strains in their capacity to sense different odorants or their *RNAi* response (Figure S1 D and E).

**Figure 1.**
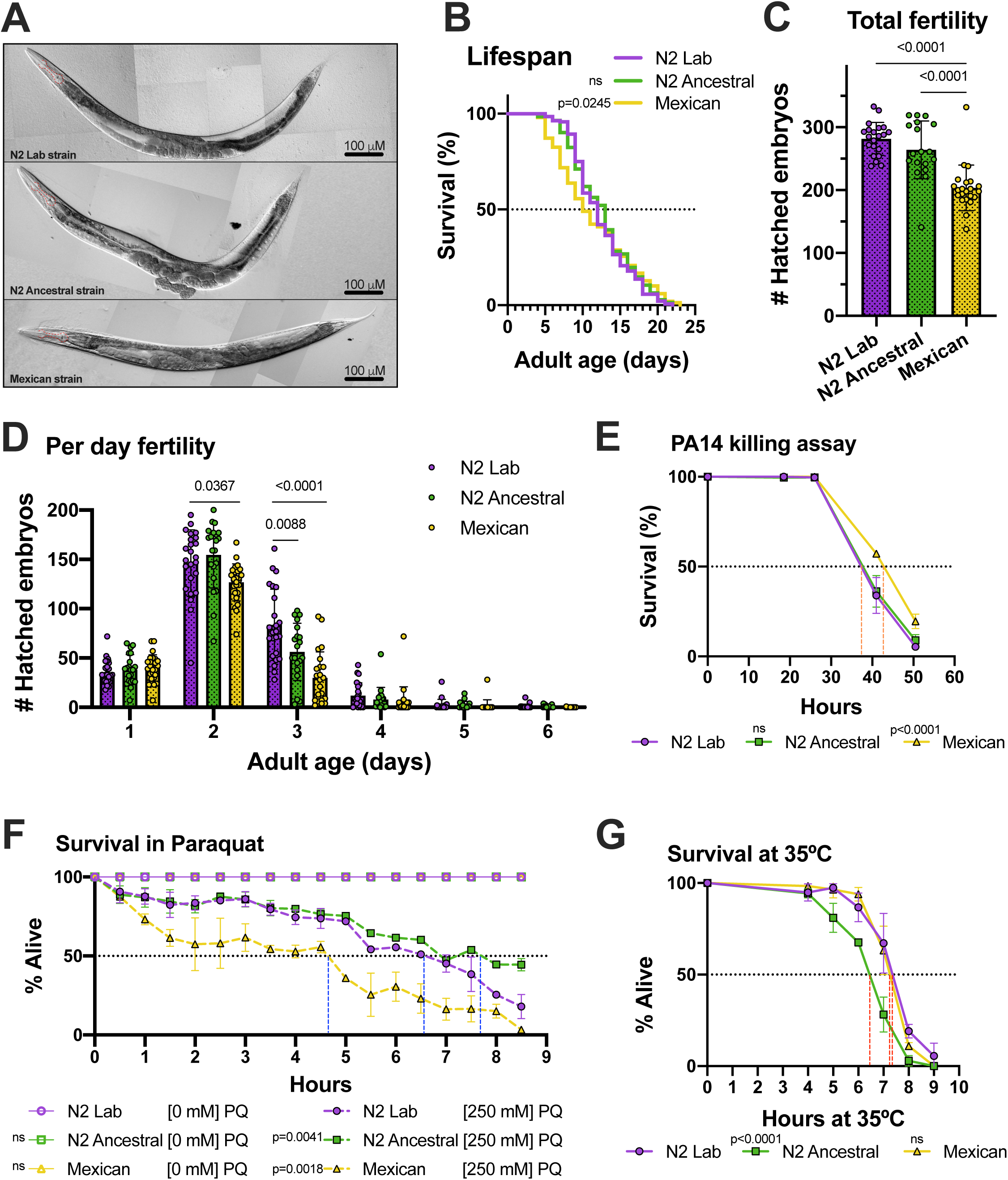
Phenotypic characterization of N2 Lab, N2 Ancestral and Mexican *C. elegans* strains. **(A)** Representative images showing similar morphology across strains. **(B)** Lifespan analysis. Data are mean values of three plates containing ∼60 worms each. Statistical significance was assessed using the Gehan-Breslow Wilcoxon test, comparing each strain to N2 Lab. Sample size: N2 Lab = 140; N2 Ancestral = 142; Mexican = 149. Median lifespans: N2 Lab=12 days; N2 Ancestral=13 days; Mexican = 10 days. **(C-D)** Total and daily fertility. Hatched embryos were counted daily for six days post-L4, one nematode per plate (day 0 = L4 stage). For total fertility all days were added. N2 Lab = 281.6; N2 Ancestral = 264.1; Mexican = 203.6. Total nematodes used: N2 Lab = 21; N2 Ancestral = 18; Mexican = 22. Significance was determined by an one-way ANOVA with multiple comparisons. **(E)** Survival curves to *Pseudomonas aeruginosa* PA14 infection. The time to 50% lethality (LT50): N2 Lab = 38 h; N2 Ancestral = 38 h; Mexican = 43 h. Three plates per strain were used, each with a PA14 lawn and ∼60 worms per plate. Hour 0 corresponds to the L4 stage. Significance was assessed by the Log-rank (Mantel-Cox) test comparing each strain to N2 Lab. **(F)** Survival curves under oxidative stress conditions. LT50: N2 Lab = 6.6 h, N2 Ancestral = 7.8 h, and Mexican = 4.7 h. ∼30 L4 worms were placed in wells containing M9 medium with food *(E. coli*) and paraquat at [250 mM] or [0 mM] for controls, with duplicates per strain. Significance was determined using the Log-rank (Mantel-Cox) test comparing each strain to N2 Lab. **(G)** Survival curves under heat stress (35 °C). LT50: N2 Lab = 7.5 h; N2 Ancestral = 6.5 h; Mexican = 7.5 h. ∼25 adult day 1 (A1) worms per plate, with three replicates per strain per hour, were exposed to a constant temperature bath at 35°C. Significance was assessed by Log-rank (Mantel-Cox) test comparing each strain to N2 Lab. All graphs show representative experiments with consistent trends across 2-3 independent replicates.

### The Mexican strain exhibits reduced lifespan and fertility under laboratory conditions

It has been reported that *C. elegans* wild isolates often exhibit lifespan differences compared to the N2 Bristol reference strain (Lee et al. 2016). Even among N2 maintained in different laboratories, variability in lifespan has been reported (Urban et al. 2021). Therefore, we evaluated the lifespan of the three strains finding that while the N2 Lab and Ancestral strains showed no statistical significant difference, having a median survival of 12 days (n = 140) and 13 days (n = 142), respectively, the Mexican strain has a shorter median lifespan of 10 days, (n = 149) (Figure 1B). To our knowledge, there are only two previous mentions regarding the lifespan of the Ancestral strain: one reported it to be shorter than the reference N2, while the other found no differences between the two (Lee et al. 2016). Our observations indicate that the tropical Mexican strain has a reduced lifespan under standard laboratory conditions.

Fertility is another key life-history trait that reflects important aspects of an organism’s biology, and reduced fertility in other wild isolates has aIso been reported (Gutteling et al. 2007; Zhang et al. 2021). We therefore evaluated both total and day-by-day fertility, finding significant differences in the three strains (Figure 1C and D). Daily fertility analysis showed that the Mexican strain had reduced offspring production on the second day compared to both N2 strains, while on the third day, both the Ancestral and Mexican strains produced fewer offspring than the N2 Lab strain (Figure 1D). The total fertility, measured as the sum of offspring produced over six days, was highest in N2 Lab (281.6, n = 21), followed by Ancestral (264.1, n = 18), and lowest in the Mexican strain (203.6, n = 22). There is a previous mention that Ancestral strain has reduced fertility in comparison to N2 Lab strain.

Interestingly, the Ancestral strain also exhibits lower fertility (at day 3) compared to the N2 Lab strain. This difference underscores the divergence between these two N2 lineages, likely reflecting evolutionary changes and adaptation to laboratory conditions. In this regard, studies on N2 domestication have shown that the laboratory-derived *nath-10* allele correlates with increased brood size by enhancing sperm production, providing a selective advantage under laboratory conditions (Duveau and Félix 2012; Sterken et al. 2015). It would be valuable to determine which *nath-10* allele is present in the Ancestral strain. On the other hand, the lower fertility observed in the Mexican strain could be attributed to multiple factors, such as differences in sperm production, oocyte quality, or embryonic viability. Also, studies have shown that changes in bacterial food sources can influence brood size and reproductive span (MacNeil et al. 2013; Stuhr and Curran 2020; Le et al. 2022). This could lead us to think that the lower fertility observed in the Mexican strain may be a consequence of exposure to the laboratory mono-bacterial diet of *E. coli* OP-50, instead of wild microorganisms.

Overall, the shorter lifespan and reduced fertility of the Mexican strain may reflect a trade-off between survival and adaptation to stable laboratory conditions.

### Strain-specific resistance profiles revealed divergent adaptation to biotic and abiotic stressors

In its natural environment, nematodes face pathogen threats. To investigate how our strains respond to biotic stress, we evaluated the survival following infection with *Pseudomonas aeruginosa* strain PA14. The Mexican strain exhibited enhanced resistance to PA14 infection, with 50% lethality occurring at 43 hour post-infection (hpi), compared to median lethal-time of 37 hpi for both the N2 Lab and Ancestral strains (Figure 1E). Previous studies have reported variation in susceptibility to bacterial infections among *C. elegans* wild isolates (Schulenburg et al. 2004; Balla et al. 2015; Martin et al. 2017). In particular, Reddy et al. (2009) showed that the wild Hawaiian isolate (CB4856) is more susceptible to PA14 infection due to the presence of the 215F allele of the neuropeptide receptor gene *npr-1*. However, Martin et al. (2017) described another Germany isolate (RC301) that also carries the same 215F allele yet exhibits enhanced resistance to PA14. These findings suggest that the Mexican strain’s resistance may involve other mechanisms, independent of the carried *npr-1* allele.

Pathogen resistance varies widely across wild isolates since significant differences in resistance to *Bacillus thuringiensis* and the microsporidian *Nematocida parissi* has been observed, indicating local adaptation to specific pathogen pressures across geographic regions (Schulenburg and Müller 2004; Balla et al. 2015). Similarly, while a Hawaiian isolate is more susceptible than N2 Lab strain to *Staphylococcus epidermidis*, it does not display increased susceptibility to PA14 (Lansdon et al. 2022). Thus it seems that variation in immune response is shaped by both geographic origin and specific pathogen encountered.

Our results suggest that Mexican strain displays a heightened immune response to PA14. This resistance could stem from genetic differences, evolutionary adaptation, transcriptional plasticity in response to a microbial-rich tropical environment, or a combination of these factors. Overall, the reduced lifespan and offspring of the tropical strain may reflect a life-history trade-off favoring enhanced resistance to pathogen threats.

In its natural environment, *C. elegans* also can encounter abiotic stressors such as oxidative stress caused by heavy metals and microorganisms (Martinez-Finley and Aschner 2011; Kniazeva and Ruvkun 2019; Mattila et al. 2022). Therefore, we exposed the different strains to paraquat (PQ) to investigate their response to reactive oxygen species (ROS) and quantify their survival to evaluate strain-specific differences in oxidative stress adaptation. Figure 1F shows that the Mexican strain was the most susceptible to PQ, with a median lethal time (LT_50_) of 4.7 hours, nearly 2 hours earlier than the N2 Lab strain (LT_50_ = 6.6 h). Interestingly, the Ancestral strain was the most resistant, with a LT_50_ of 7.8 hours. The Mexican strain carries the “social” *npr-1* 215F allele, which is associated with differences in oxygen tolerance (Gray et al. 2004; Sterken et al. 2015; Zhao et al., 2018), potentially contributing to its reduced PQ tolerance. However, it is less likely that the differences in ROS sensitivity between the N2 Lab and Ancestral strains are explained by variation in the *npr-1* allele, as both presumably carry the same version. However, another domestication-related allele, *glb-5,* has also been reported to influence responses to oxygen levels and was present in various strains prior to the original N2 Bristol cryopreservation (McGrath et al. 2009; Persson et al. 2009; Sterken et al. 2015). Therefore, it is possible that the fixation of *glb-5* allele occurred after the divergence of the N2 Lab and Ancestral strains, but such hypothesis requires further validation.

Another abiotic stressor we evaluated was temperature, particularly considering the weather differences between the geographical origin of the strains. We exposed adult worms to 35 °C and monitored their survival every hour (see methods). The Mexican and N2 Lab strains exhibited similar survival profiles, with a lethal median time (LT_50_) of 7.5 h, whereas the Ancestral strain was more susceptible to heat stress with a LT50 of 6.5 hours (Figure 1G). Mexican strain comes from a tropical environment, where temperatures range from 7 °C to the 35.4 °C, highest registered temperature in Mexico City; while Bristol (UK), origin of the N2 strain, experiences a colder climate, ranging from 2.8 °C to 20 °C. The Ancestral strain may still retain adaptations to cooler environments, which could explain its increased sensitivity to elevated temperatures. Another hypothesis is that Ancestral strain’s enhanced resistance to ROS may come at the cost of reduced thermotolerance, given the interplay between heat stress and oxidative stress responses (Crombie et al. 2016). It would also be possible to hypothesize that the N2 Lab strain has undergone adaptation to a warmer and more stable laboratory environment of 20 °C, which could account for its heat stress resistance comparable to that of the tropical strain. It would be informative to assess the survival of both the N2 Lab and the Mexican strain under less acute heat stress conditions to better understand how long-term maintenance at a constant 20 °C may shape thermal tolerance.

Together, these results reveal that each strain exhibit distinct stress resistance profiles: The Mexican tropical strain excels in pathogen resistance but is more vulnerable to oxidative stress; the Ancestral strain shows heightened oxidative stress tolerance but reduced thermotolerance and pathogen resistance; and the N2 Lab strain shows adequate oxidative stress and temperature tolerance.

### Transcriptomic profiles reveal greater similarity between wild isolates

The observed phenotypic variability of life-traits among the three strains may arise not only from genetic but also transcriptional variability. To explore this, we analyzed their transcriptomic profiles by RNA-seq under standard laboratory conditions. Principal component analysis (PCA) showed that each strain replicate clustered tightly, indicating high intra-strain consistency. However, the N2 Lab strain clustered distinctly from the other strains along the first principal component (PC1), which accounted for 78% of the total variance. In contrast, the Mexican and Ancestral strains were separated primarily along the second principal component (PC2), explaining 13% of the variance (Figure 2A). We assessed the global expression level of the transcriptome quantified by transcript per million (TPM) finding statistical differences between all the strains (Figure 2B). These results suggest greater transcriptional divergence between the N2 Lab strain and the wild isolates than between the wild isolates themselves. These observations are consistent with previous findings that the Bristol N2 strain cluster apart from other wild isolates, showing strain-specific transcriptomic profiles (Lansdon et al. 2022).

**Figure 2.**
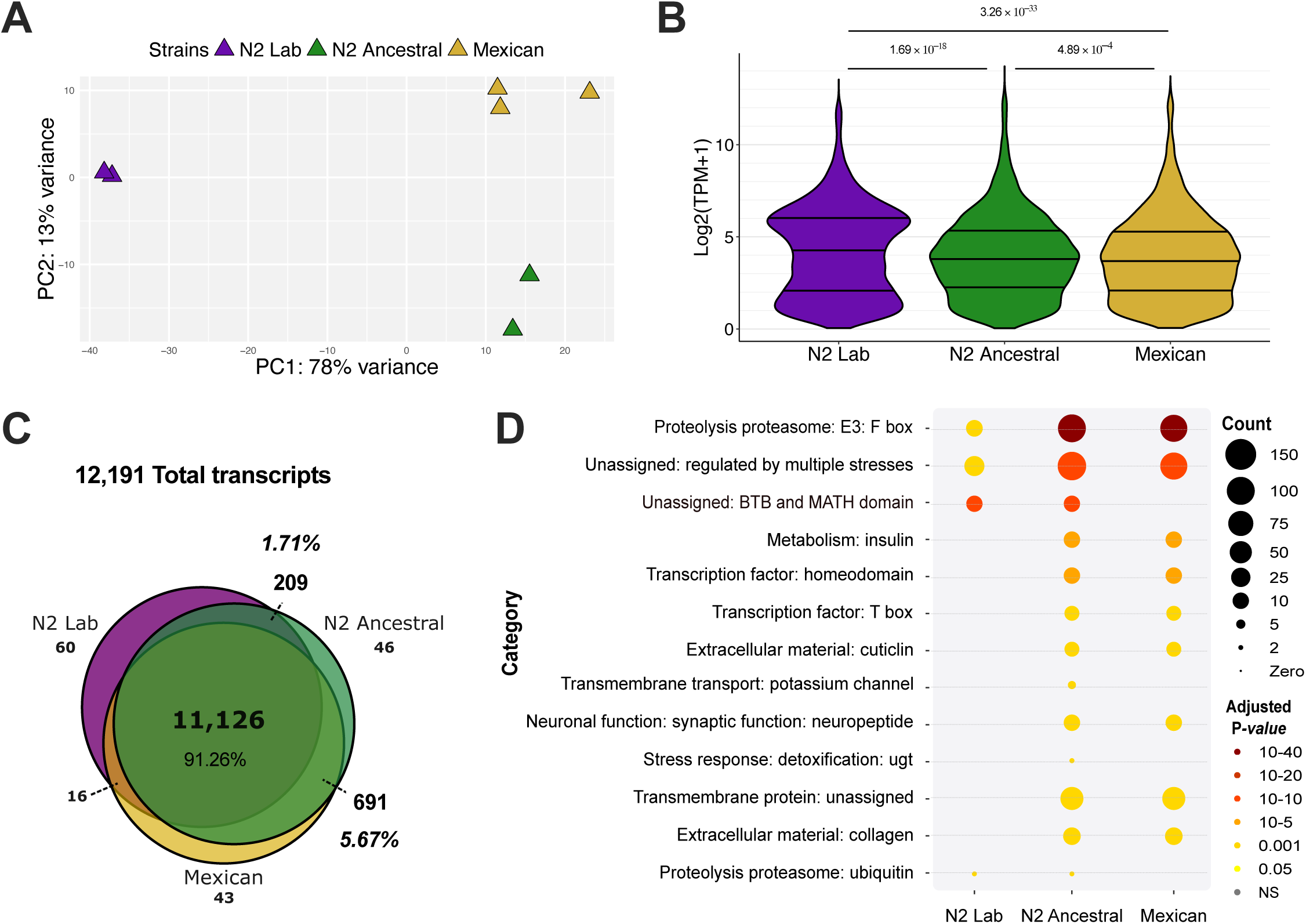
Strain-specific gene expression profiles. **(A)** Principal component analysis (PCA) shows clustering of samples by strain. **(B)** Overall transcript expression pattern, measured in Transcripts Per Million (TPM). Kruskal-Wallis test showed significant differences (*x^2^ = 154.87, p = 2.2 × 10^−16^*), Dunn post-hoc test, with *P-values* adjusted by Benjamini-Hochberg, showed significant differences between paired comparisons (*P-adjusted* shown on plot). **(C)** Venn diagram showing that 11,126 genes (91.26% of all expressed genes) arre shared among the three strains. **(D)** Enriched Gene Ontology Terms associated with groups of expressed transcripts: N2 Lab-Ancestral, Ancestral-only, Ancestral-Mexican, and Mexican-only. All terms have a FDR-adjusted *P*-value < 0.01 as a measure of enrichment (Fisher exact test; Bonferroni correction) of the GO term among expressed genes; WormCat category = 3 (See Table S1).

To assess the extent of shared expression, we compared the expressed transcript set across all strains. We identified a common core universe composed of 11,126 transcripts (91.26% of the total 12,191), shared by all strains (Figure 2C). These transcripts likely correspond to the core genes of adult day 1 worms growing in standard laboratory conditions. Interestingly, the Ancestral strain shared more expressed transcripts with the Mexican strain (691 = 5.67%) than with the N2 Lab strain (209 = 1.71%) (Figure 2C), highlighting a stronger transcriptomic affinity between wild isolates. We next performed gene set enrichment analysis of each of the six transcript groups: N2 Lab-only, N2 Lab-Ancestral, Ancestral-only, Ancestral-Mexican, Mexican-only, and Mexican-N2 Lab. Neither of the 60 N2 Lab-only transcripts nor the 16 shared exclusively between Mexican-N2 Lab strains yielded any significant Gene Ontology (GO) terms (FDR-adjusted *P-value* < 0.01) (Table S1). Among the remaining comparisons, the GO term “Proteolysis proteasome: E3: F-box” was consistently enriched in all of them (Figure 2D, Table S1), which relates to the ubiquitin mediated protein degradation pathway, responsible for maintenance of the normal cellular physiology (Ghazi et al. 2007; Papaevgeniou and Chondrogianni 2014). This GO category has been previously reported as highly enriched when analyzing gene expression variation among wild isolates (Zhang et al. 2022).

Among the 46 Ancestral-only genes, we observed two additional categories beyond protein degradation: “Transmembrane transport: potassium channels” and “Stress response: detoxification: UGT”. Potassium channels, encoded by ∼70 genes in *C. elegans*, are expressed across multiple cell types, including neurons (Salkoff et al. 2005) while UDP Glycosyltransferases (UGTs) are enzymes that mediate clearance of toxic compounds by glucuronidation. *C. elegans* contains more than 70 *ugt* genes, in contrast with only 22 in humans (Asif et al. 2024; Gómez-Orte et al. 2017). Interestingly, *ugt* genes have been previously reported to be up-regulated in worms fed with *E. coli* compared to those fed with *B. subtilis*, due to toxic redox activity associated with coenzyme Q biosynthesis (Gómez-Orte et al. 2017).

On the other hand, the Ancestral-N2 Lab group (209 transcripts) was enriched in “Proteolysis proteasome”, the “BTB and MATH domain” and “Unassigned: regulated by multiple stresses”. In contrast, the Ancestral-Mexican group (691 transcripts), exhibited a broader range of GO categories, including “Unassigned: regulated by multiple stresses”, “Transmembrane protein”, “Metabolism: insulin”, “Transcription factor”, “Extracellular material” and “Neuronal function” GO terms (Figure 2D, Table S1), suggesting coordinated regulation of physiological and stress-related processes shared between these wild isolates.

Overall, these results highlight a greater number and diversity of significantly enriched GO terms shared between the Ancestral and Mexican strains, further supporting their closer transcriptomic relationship compared to either with the N2 Lab strain (Table S1).

### Differential gene expression landscape reveal specific wild strains patterns

As PCA and overall transcriptional profiles suggest that the Mexican and Ancestral strains are more similar to each other than to the lab-domesticated strain, we performed differential gene expression (DEG) analyses comparing both Mexican and Ancestral strains against N2 Lab, using a cut-off of FDR-adjusted p-value < 0.05 and log2 fold change (LFC) ≥ 2. This revealed a total of 3,400 DEGs across both comparisons, with 55% (1,873) shared by the two wild strains (611 downregulated, 1,262 upregulated) (Figure 3A-B). Heatmap analysis clearly shows that the expression profiles of the two wild strains are markedly different from that of the N2 Lab strain, with DEGs representing approximately 28% of all expressed genes (3,400/12,191) (Figure 3C). This extensive transcriptional divergence between wild and lab strains aligns with previous reports highlighting significant gene expression shifts in wild isolates and in response to environmental factors such as diet (MacNeil et al. 2013; Snoek et al. 2017; Zhang et al. 2022). We also examined known regulators of transcriptional variability in wild isolates. Both *hil-2* (a linker histone cofactor) and *ttx-1* (a thermosensory transcription factor), previously implicated in gene expression differences among wild strains, were upregulated in Mexican and Ancestral strains (Table S2) contributing to these wild strains different expression profiles (Zhang et al. 2022).

**Figure 3.**
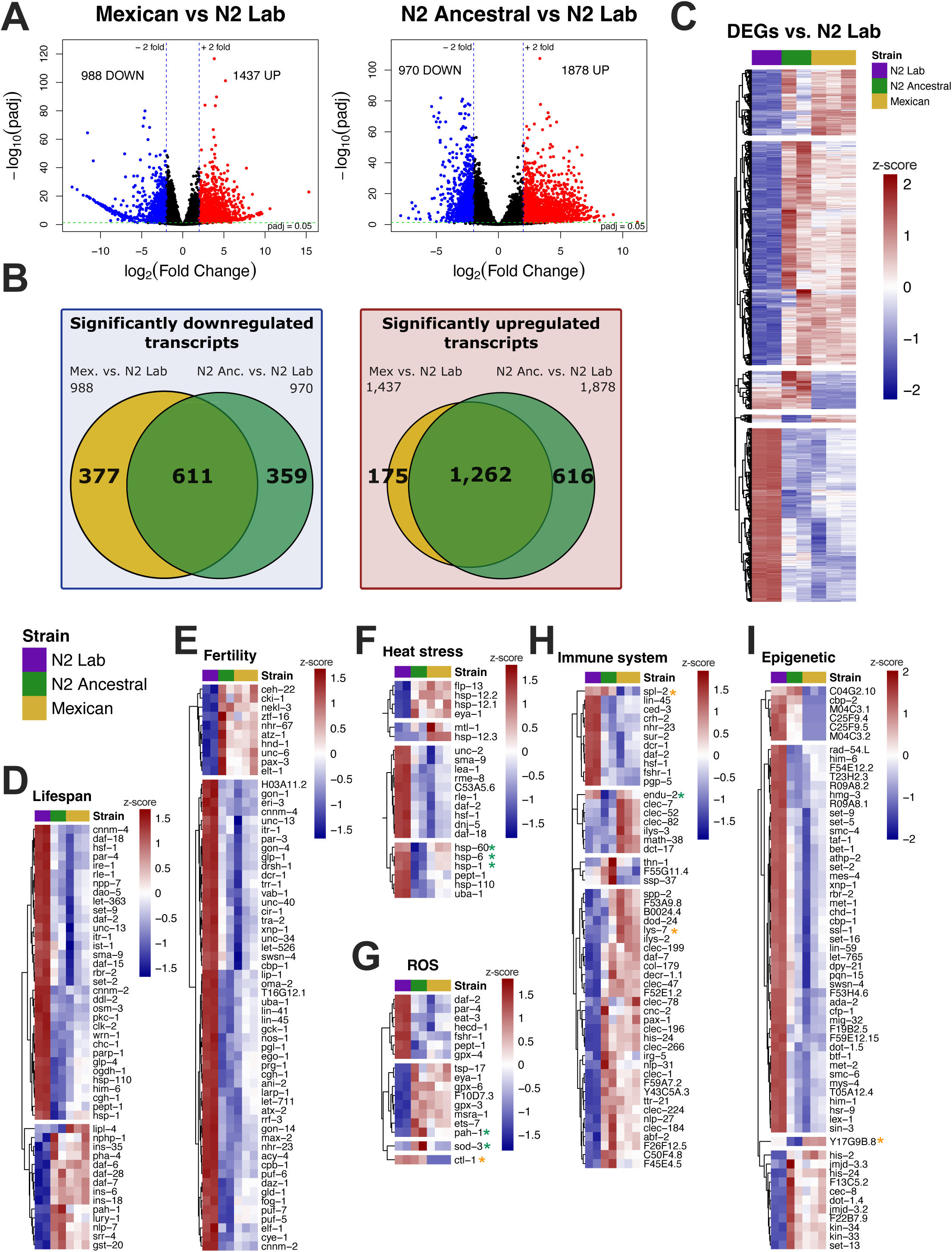
Differential gene expression relative to the N2 Lab strain. **(A)** Volcano plots showing differentially expressed genes (DEGs) of Mexican and Ancestral strains compared to N2 Lab strain. DEGs were defined as those with FDR-adjusted *P-value* < 0.05; log_2_ fold change (LFC) ≥ 2. **(B)** Venn diagrams showing the overlap of up-regulated and down-regulated transcripts relative to the N2 Lab strain. Mexican and N2 Ancestral strains share 61.5% (1,262/2053) of up-regulated and 45.4% (611/1347) of down-regulated transcripts. **(C)** Heatmap for the total 3,400 DEGs vs. N2 Lab strain. **(D-I)** Heatmaps for genes grouped by functional categories: *Lifespan*, *Fertility*, *Heat stress*, *ROS*, *Immune system* and *Epigenetic regulation*. All DEGs have a *P*-adjusted value ≤ 0.05 and a LFC ≥ 2. Gene group annotations were obtained from GO terms using AmiGO (see Methods).

### Upregulated genes relate to immune system and pathogen response

Venn diagram-based grouping of DEGs allowed us to identify strain-specific and shared expression patterns; Gene Ontology (GO) enrichment analyses were then conducted for each group (Table 1). Among Mexican-only upregulated genes, GO terms such as “Stress response: pathogen” and “Proteolysis general: lysozyme” emerged, consistent with this strain’s enhanced resistance to PA14 (Figure 1E). After building a heatmap on immune system related genes, we found upregulation of *lys-7* (Figure 3H), a lysozyme gene downregulated by PA14 during infection as a virulence strategy, that when its expression gets restored, *C. elegans* is rescued from mortality, highlighting its importance in defense against this pathogen (Evans et al. 2008; Fatin et al. 2017). Conversely, Mexican-only downregulated genes included *spl-2*, a sphingosine phosphate lyase whose hyperactivation increases susceptibility to PA14 (Nasrallah et al. 2023), further aligning with the enhanced immunity of this strain. The enrichment of immune-related GO terms among Mexican-specific upregulated genes underscores the link between transcriptional changes and the strain’s superior pathogen resistance (Figure 1E).

**Table 1.**
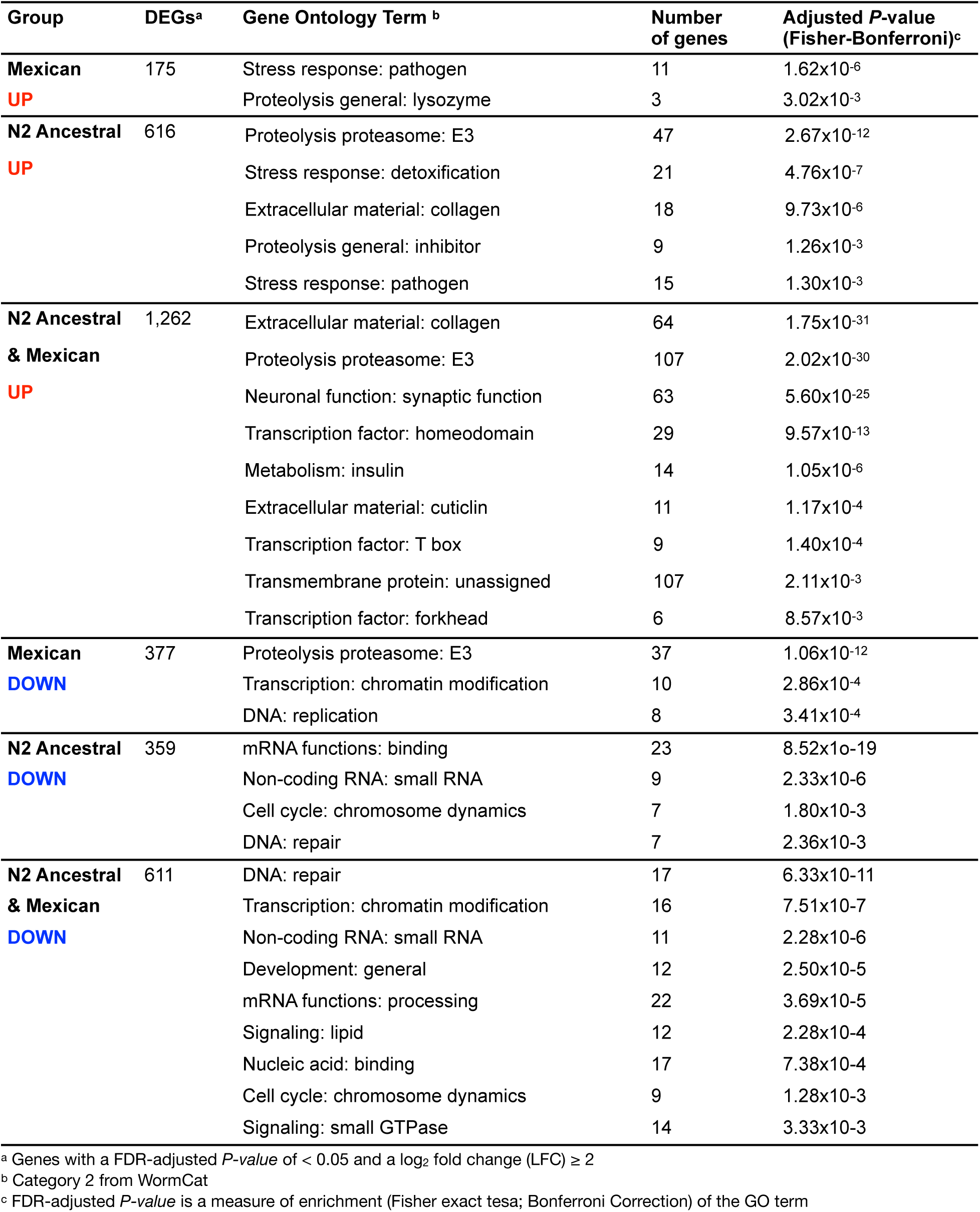
Enriched Gene Ontology Terms for differentially expressed genes (DEGs) of N2 Ancestral vs. N2 Lab strain, and Mexican vs. N2 Lab strain.

Within Ancestral-only upregulated genes, terms such as “Proteolysis proteasome: E3”, “Stress response: detoxification”, “Extracellular material: collagen”, and “Stress response: pathogen” were shown, which may relate to pathogen resistance or environmental adaptation (Table 1). Additionally, three Ancestral-specific immune genes appeared upregulated in the immune-related heatmap (third cluster, figure 3H), highlighting how distinct pathogen exposure histories could shape differential immune strategies among wild isolates (Schulenburg and Müller 2004; Balla et al. 2015). Such variation in immune-related gene expression may reflect the evolutionary impact of unique environmental pressures on each wild isolate.

The shared DEGs between Mexican and Ancestral strains included upregulation of collagen genes and terms such as “Neuronal function: synaptic function” (Table 1). The upregulation of collagen genes may be linked to environmental adaptation, as the collagen rich cuticle acts as a protective barrier. Over 60 collagen transcripts were upregulated in both wild strains (Table 1), likely due to upregulation of the transcription factor *elt-3* in both strains (Table S2), which regulates collagen gene expression in response to environmental stress (Mesbahi et al. 2020). Considering that *C. elegans* possesses 173 predicted cuticular collagen genes (Teuscher et al. 2019), such extensive collagen gene upregulation likely reflects an adaptive response to environmental stresses encountered by wild strains.

### Shared Mexican-Ancestral downregulated genes associate with chromatin remodeling

Interestingly, Ancestral and Mexican downregulated genes were enriched in “DNA: repair”, “Non-coding RNA: small RNA”, “Nucleic acid: binding”, and “Transcription: chromatin modification” (Table 1). This prompted a closer examination of epigenetic regulators (after building a heatmap) where we identified six chromatin remodeling genes downregulated exclusively in the Mexican strain, alongside *Y17G9B.8*, upregulated in the Mexican strain and downregulated in the Ancestral strain (Figure 3I), which encodes the ortholog of human SGF29, a chromatin reader involved in histone H3 acetylation (Bian et al. 2011; Sternberg et al. 2024), suggesting that chromatin remodeling factors may contribute to strain-specific transcriptional regulation.

### Differential expression of heat and oxidative stress genes links to phenotypic variation

To contextualize transcriptional differences in terms of phenotypic outcomes, we explored DEGs associated with known life-history traits such as fertility, lifespan, and stress responses. Only one lifespan-related gene, *lipl-4*, was found upregulated exclusively in the Mexican strain, although this did not correspond to lifespan extension (Figure 3D). Fertility-related DEGs showed similar expression in both wild strains (Figure 3E). While *lipl-4* upregulation is generally associated with lifespan extension (Folick et al. 2015), its lack of effect here suggests additional factors modulate lifespan in this strain.

We also examined heat- and oxidative stress-related genes. The Ancestral strain showed downregulation of heat shock genes including *endu-2*, *hsp-1*, *hsp-6*, and *hsp-60*, which are associated with reduced thermotolerance (Figure 3H and 3F). In particular, downregulation of *endu-2*, that is implicated in transcriptional reprogramming following heat stress (Xu et al. 2023), may contribute to the observed reduced heat tolerance. Remarkably, using a lower threshold (LFC ≥ 1.5), revealed additional downregulated heat shock genes (*hsp-90*, *cct-2*, *cct-4*, *cct-8*, *rpn-2*, *uba-1*) involved in canonical heat shock response (Table S2; Guisbert et al. 2013). Regarding oxidative stress response, the Mexican strain (the most ROS-sensitive; Figure 1F), has uniquely downregulated *ctl-1*, a catalase critical for H₂O₂ detoxification (Schiffer et al. 2020; Branicky et al. 2022). In contrast, the Ancestral strain, witch exhibited the highest paraquat resistance, showed upregulation of *sod-3*, *pah-1* (Figure 3G), and *gst-4*, *gst-10* and *gpx-5* (with a lower threshold LFC ≥ 1.5; Table S2), genes linked to antioxidant defenses (Ayyadevara et al. 2005; Calvo et al. 2008; Van Raamsdonk and Hekimi 2012). Together, these findings underscore strain-specific strategies for coping with environmental stress.

Finally, we directly compared the gene expression of Mexican and Ancestral strains showing that only 6.8% of the DEGs (127 down- and 51 up-regulated, out of 2,603 total DEGs) were specific between these two strains (Figure 4), further supporting transcriptional similarity between wild-isolates. GO analysis of these DEGs revealed only a few significant terms, including “Stress response: detoxification” and “Stress response: pathogen” among Mexican vs. Ancestral downregulated genes, reflecting subtle strain-specific adaptations (Table 2). One notable GO term was “Major sperm protein,” downregulated in the Mexican strain and consistent with its reduced fertility (Table 2, Figure 1C-D). These subtle expression differences likely contribute to nuanced phenotypic variation between the two wild strains, reinforcing the close but distinct transcriptional profiles observed.

**Figure 4.**
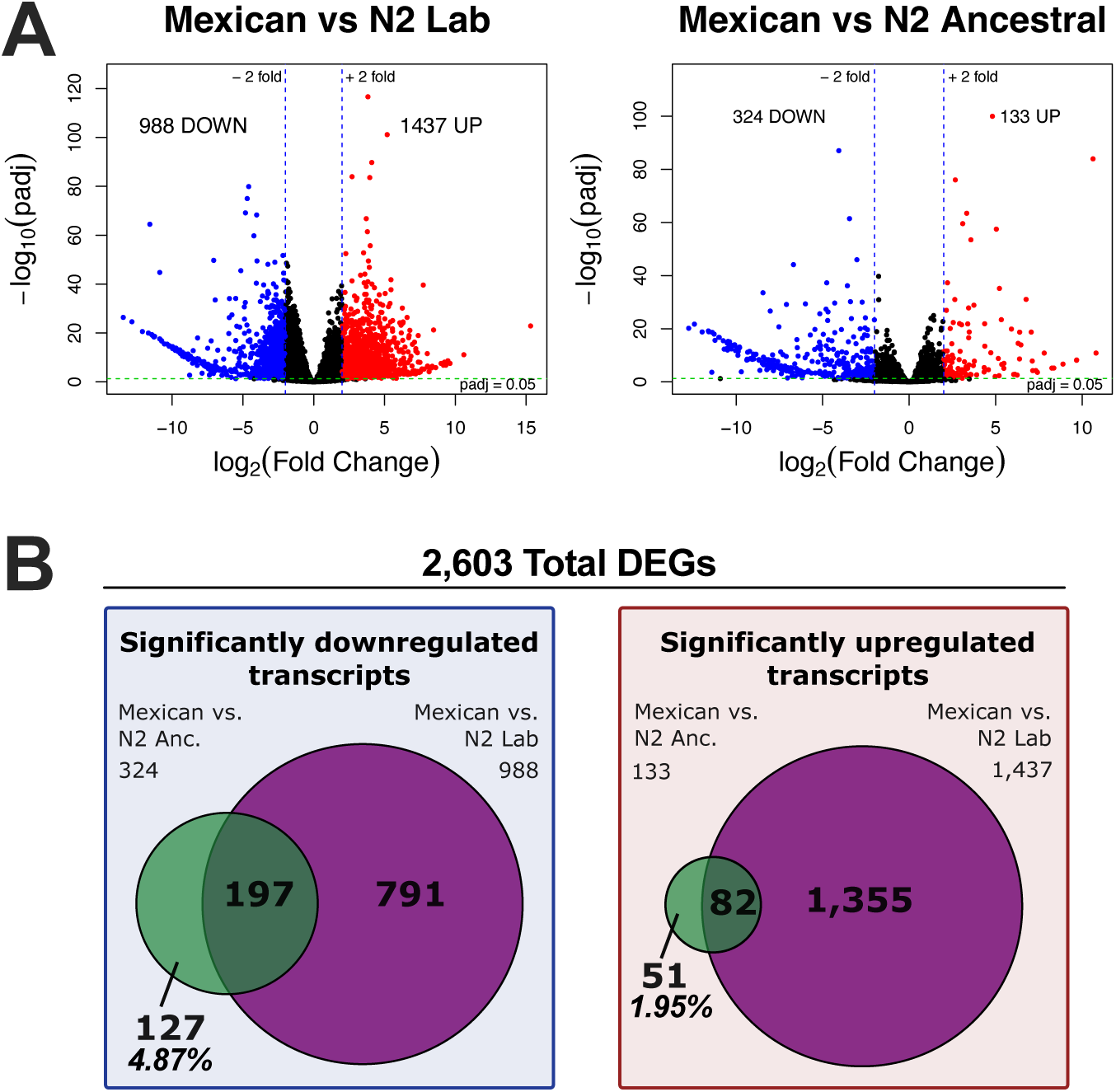
Differentially Expressed Genes in the Mexican strain compared to N2 Lab and Ancestral strains. **(A)** Volcano plots showing differentially expressed genes (DEGs) of Mexican vs. N2 Lab, and Mexican vs. Ancestral strains comparisons. FDR-adjusted *P-value* < 0.05; log_2_ fold change (LFC) ≥ 2. **(B)** Venn diagrams showing shared up- and down-regulated transcripts of Mexican strain vs. N2 Lab and Ancestral strains. N2 Lab and Ancestral strains share 5.5% (82/1488) up-regulated and 17.7% (197/1115) down-regulated transcripts.

**Table 2.**
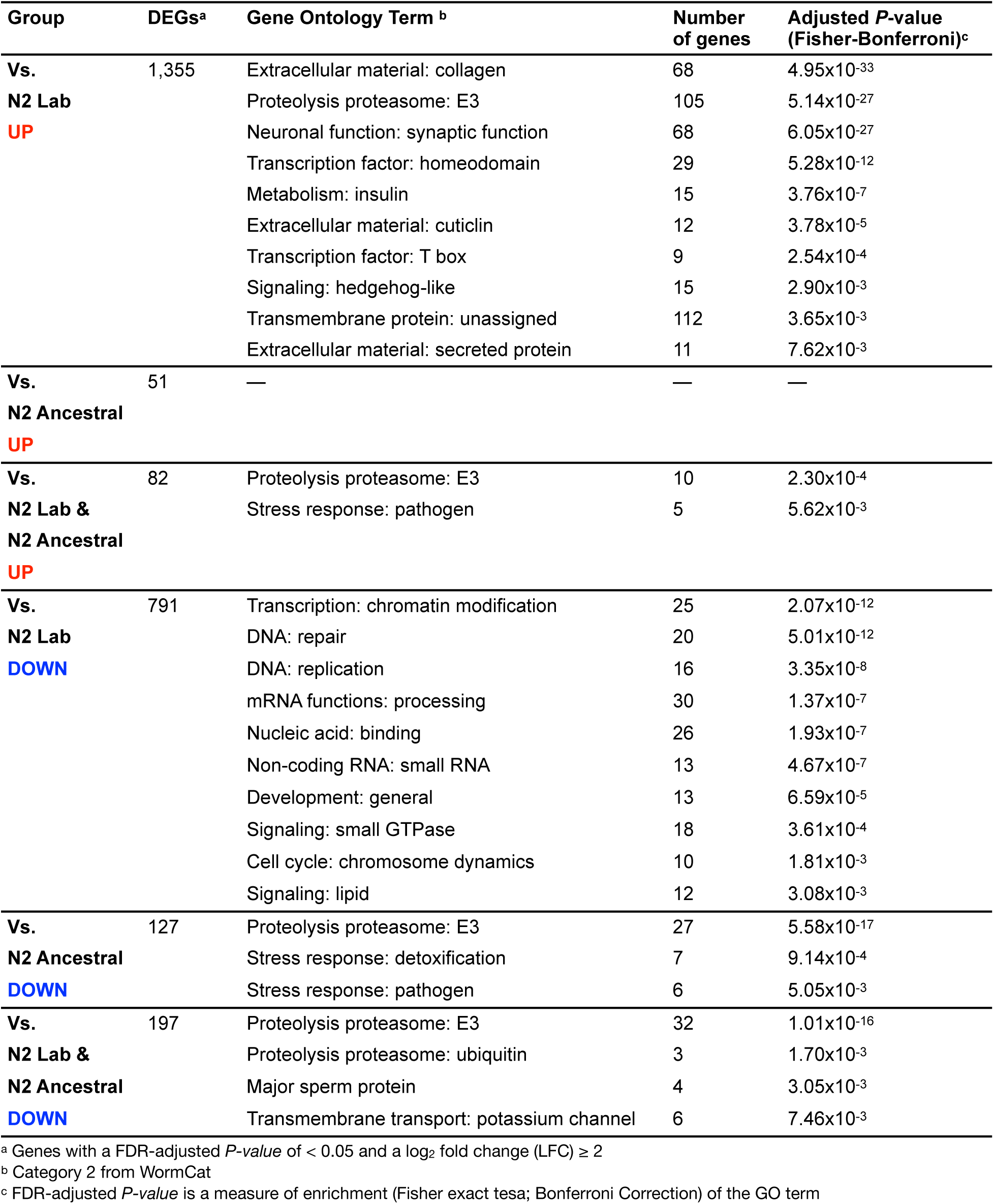
Enriched Gene Ontology Terms for differentially expressed genes (DEGs) of Mexican vs. N2 Lab strain, and Mexican vs. N2 Ancestral strain.

Overall, these findings underscore the complexity of regulatory networks driving transcriptional divergence between lab-domesticated and wild *C. elegans* strains.

### Gene expression patterns under acute heat stress revealed both conserved and strain-specific thermal adaptive transcriptional responses

Thermal adaptation is essential for organism survival; therefore we investigated gene expression differences in response to heat stress, particularly considering the increased temperature sensitivity of the Ancestral strain (Figure 1G). To this end, we exposed the animals to a sublethal heat stress protocol (2 hours at 35 °C) and generated a new set of RNA-seq for all the strains. PCA revealed that replicates from each strain cluster together and, again, the N2 Lab strain diverged greatly along the principal component 1 (PC1), accounting for 93% of the variance, whereas the Ancestral and Mexican strains separated primary on the second axis (PC2) by 5% variance (Figure 5A). Global expression levels measured by TPM further supported this pattern, showing closer resemblance between the Mexican and Ancestral strains (Figure 5B). Of the 12,356 expressed transcripts, 84.9% were shared among all strains under heat stress (Figure 5C), with gene ontology analysis revealing enrichment in categories such as stress response, proteolysis, cytoskeleton and transcription factors (Figure 5D, Table S3), consistent with a generalized response to high temperature.

**Figure 5.**
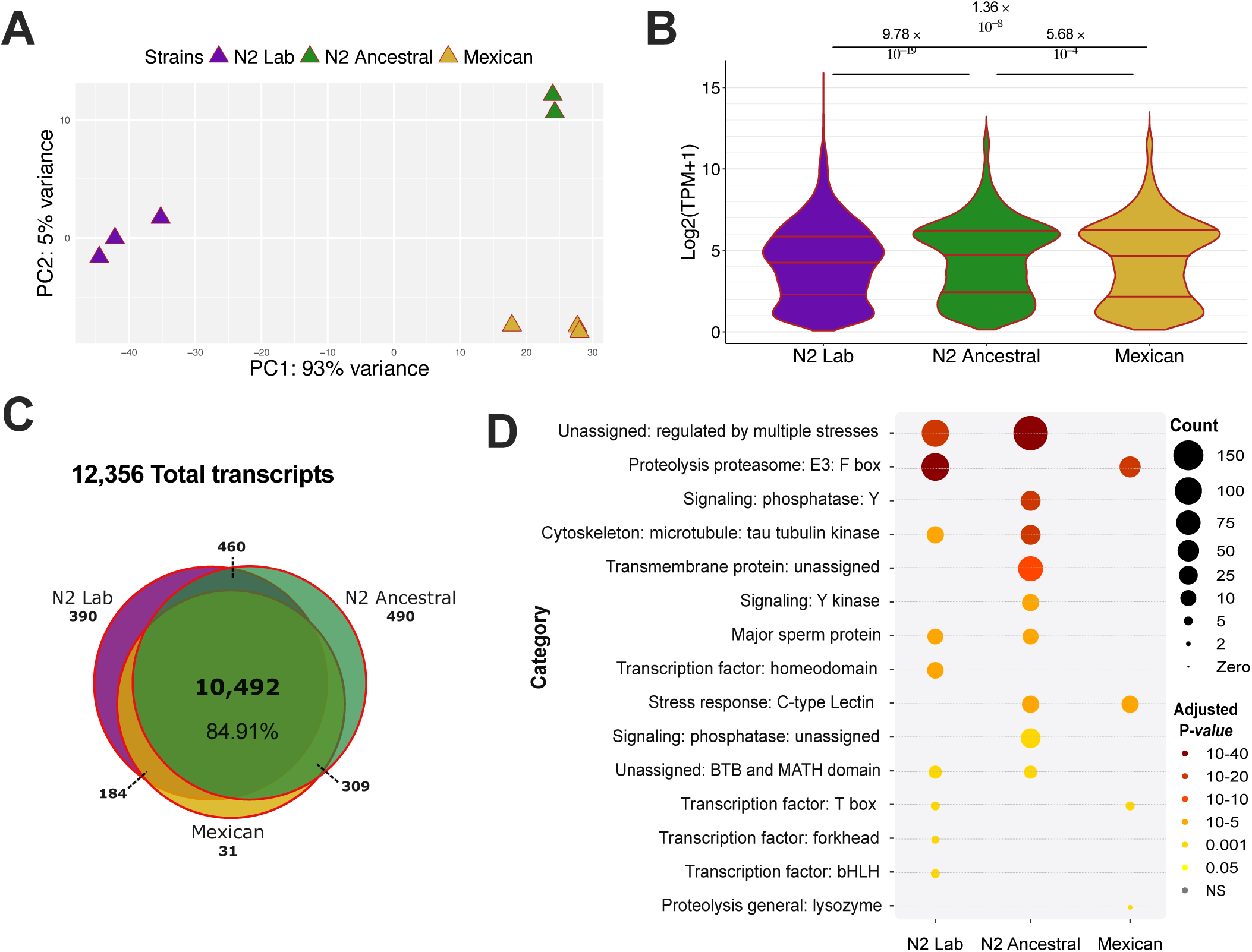
Strain-specific gene expression profiles under heat stress (35 °C) **(A)** Principal component analysis (PCA) shows clustering of samples by strain. **(B)** Overall transcript expression pattern, measured in Transcripts Per million (TPM), differs in all strains. Krustal-Wallis test showed significant differences (*x^2^ = 80.78, p = 2.2 × 10^−16^*), Dunn post-hoc test with *P-values* adjusted by Benjamini-Hochberg showed significant differences between paired comparisons (*P-adjusted* showed on plot). **(C)** Venn diagram showing that 10,492 genes (84.91% of all expressed genes) are shared across the three strains at 35 °C. **(D)** Enriched Gene Ontology Terms for expressed genes at 35°C. Terms corresponding to the transcripts groups: N2 Lab-only, N2 Lab-Ancestral, Ancestral-only, Ancestral-Mexican, Mexican-only, and Mexican-N2 Lab. All terms have a FDR-adjusted *P*-value < 0.01 as a measure of enrichment (Fisher exact test; Bonferroni correction) of the GO term among expressed genes; WormCat category = 2 (See Table S3).

To identify DEGs, we compared transcriptomes at 35 °C and 20 °C within each strain. All three strains mounted robust transcriptional responses to heat stress, with 2,486 DEGs in the N2 Lab strain, 2,942 DEGs in the Ancestral strain and 2,944 DEGs in the Mexican strain (Figure 6A) with characteristic overexpression of heat shock proteins (HSPs), as evident in a distinct HSP cluster (Figure 6C). Surprisingly, only 0.91% (27/2963) of the total up-regulated DEGs and 1.6% (54/3356) of total down-regulated DEGs, were shared across all three strains under heat stress (Figure 6B). GO analysis among the 27 shared up-regulated DEGs identified only one significant term: “Stress response: heat”, driven by strong induction (∼8 LFC) of three genes: *hsp-16.2*, *hsp-16.41* and *hsp-70* in all strains (Table 3, Table S2), underscoring their central role in canonical heat stress response. On the other hand, the Mexican and Ancestral strains shared a substantial large proportion of DEGs: 23.92% (709/2963) of up-regulated and 32.59% (1094/3356) of down-regulated (Figure 6B), highlighting their transcriptional similarity under stress conditions. GO analysis of these shared DEGs revealed enrichment in terms related to “DNA: repair”, “Transcription” and “Signaling”, for up-regulated, and to “Neuronal function: synaptic function”, “Transmembrane protein” and “Proteolysis proteasome: E3”, for down-regulated (Table 3).

**Figure 6.**
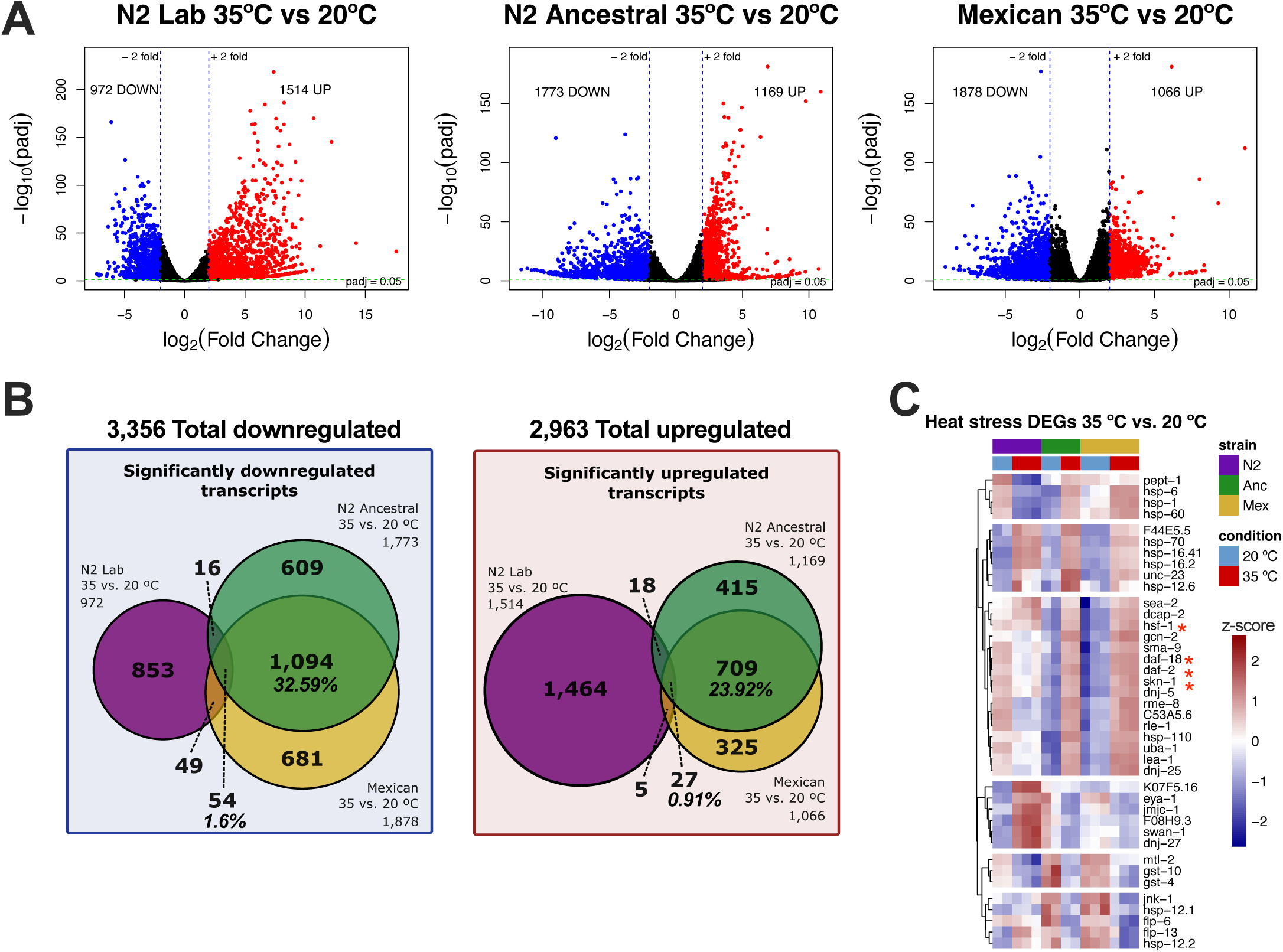
Differentially Expressed Genes in each strain under heat stress. **(A)** Volcano plots showing differentially expressed genes (DEGs) at 35 °C compared to 20 °C for each strain. FDR-adjusted *P-value* < 0.05; log_2_ fold change (LFC) ≥ 2. **(B)** Venn diagrams showing shared up- and down-regulated transcripts of the three strains at 35 °C. Shared by the three strains= 0.91% (27/2963) up-regulated; and 1.6% (54/3356) down-regulated **(C)** Heatmap showing DEGs associated with the to *Heat stress* GO term. All DEGs have a *P*-adjusted value ≤ 0.05 and a LFC ≥ 2. Gene group list was recovered by GO terms using Gene Ontology tool AmiGO (see Methods).

**Table 3.**
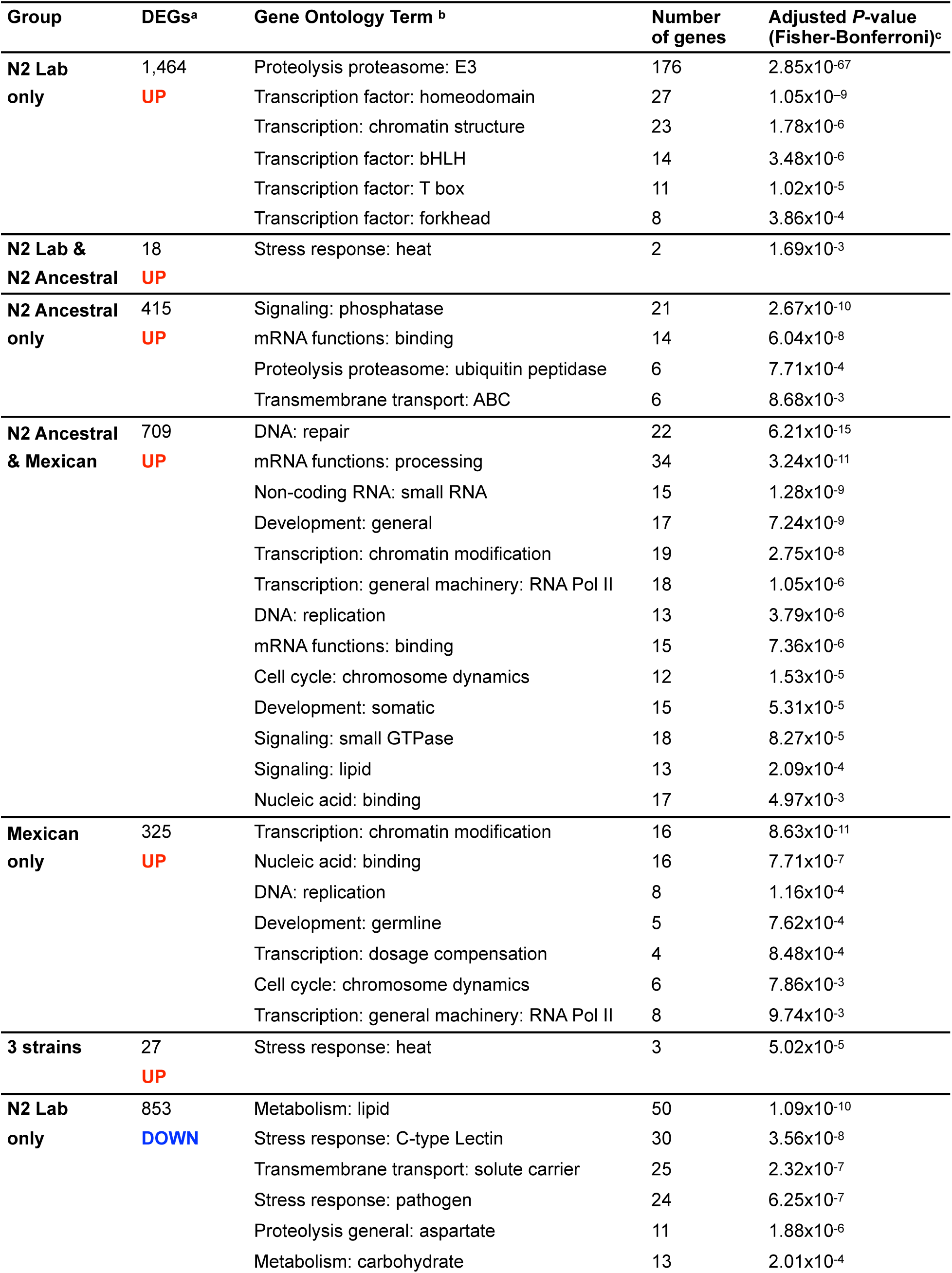
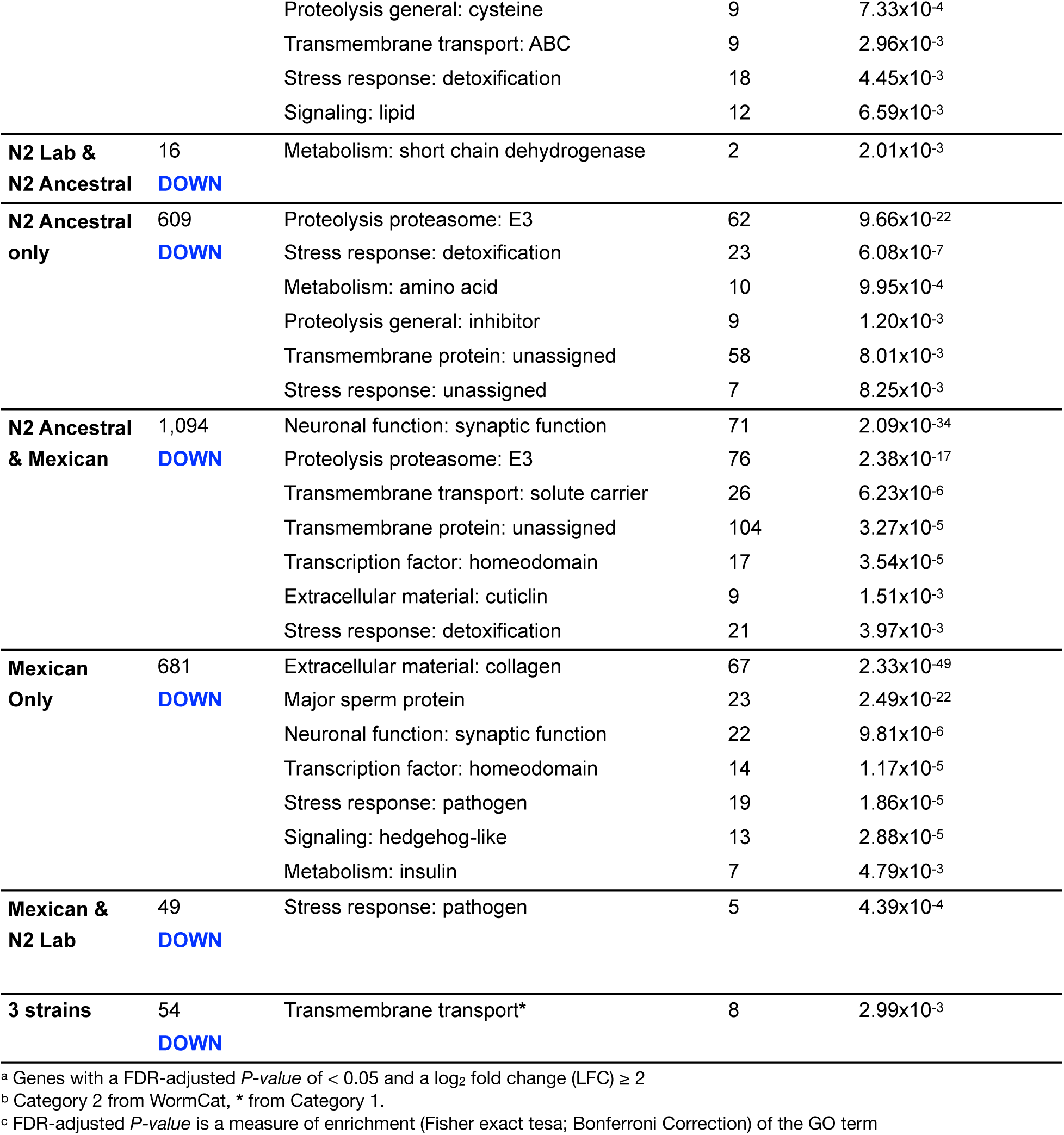
Enriched Gene Ontology Terms for differentially expressed genes (DEGs) 35 °C vs. 20 °C.

To further dissect strain-specific transcriptional dynamics, we analyzed DEGs that were grouped by Venn diagrams (Figure 6B). Among the few shared transcripts upregulated in both the Ancestral and N2 Lab strains (18 genes), only two were heat-stress related: *hsp-12.6* and *F44E5.5* (Figure 6B, Table 3, Table S2). *F44E5.5* encodes an ortholog of human HSPA6/HSPA7, and in *C. elegans* is upregulated by the heat shock transcription factor HSF-1 (Brunquell et al., 2016). Although *hsp-12.6* and *F44E5.5* were not identified as DEGs in the Mexican strain, *hsp-12.6* expression was detected at almost significant levels (LFC: 1.22, adjusted p-value: 0.054), suggesting its involvement in the heat response.

In the Ancestral strain, upregulation of *hsp-6* and *hsp-60* stood out as part of a cluster that also includes *hsp-1*. This same cluster showed moderate upregulation in the Mexican strain but was downregulated in the N2 Lab strain (first cluster, Figure 6C, Table S2). While *hsp-1* is directly related to HSR, *hsp-6* and *hsp-60* are involved in mitochondrial unfolded protein response, and *pept-1,* another gene in this cluster, encodes a transmembrane transporter (Sternberg et al. 2024). These patterns suggest that mitochondrial stress pathways may be more strongly activated in the wild isolates.

Notably, among the 325 Mexican-specific upregulated DEGs (Figure 6B), *hsf-1* was the only canonical heat shock response (HSR) gene identified. HSF-1 regulates proteostasis under heat stress by inducing chaperones such as as *hsp-16.1*, *hsp-16.2*, *hsp-16.41* and *hsp-16.48,* promoting protein refolding and degradation of damage ones (van Oosten-Hawle and Morimoto 2014; Jones et al. 2018). Interestingly, *hsf-1* clusters with other 15 genes, upregulated in the Mexican and Ancestral strains, including *dnj-5*, *rme-8*, *hsp-110, daf-2*, *daf-18,* and *skn-1* (third cluster, Figure 6C, Table S2 and Figure S2); genes involved in stress signalling and longevity pathways (Jones et al. 2018; Zhang et al. 2001; Sternberg et al. 2024). Notably, no changes were detected on these genes in the N2 Lab strain, but their expression levels were higher at 20 °C, supporting the existence of a more dynamic response program in the wild isolates.

Finally, the N2 Lab-specific cluster of upregulated genes included transcripts like *swan-1* (involved in heat/osmotic response; Ritterhoff et al. 2010)*, eya-1* (which down-regulation decreases worm’s heat and oxidative stress tolerance; Wang et al. 2014); the transcriptional factor *jmjc-1* (regulator of stress associated genes; Kirienko and Fay 2010), and *F08H9.3*, (associated to desiccation resistance; Erkut et al. 2013). These findings suggest a distinct, perhaps lab-adapted, stress response in the N2 Lab strain, potentially shaped by its long-term cultivation.

Overall, these results reveal a complex interplay of conserved and strain-specific gene regulatory programs that underlie the thermal adaptive response in *C. elegans*. Heat stress response (HSR) at 2 hour-exposure is more alike in Mexican and Ancestral strains, perhaps as a consequence of different transcriptional dynamics over heat exposure versus N2 Bristol (Jovic et al. 2017), knowing that at longer exposure times, they behave differently.

## Discussion

Animal adaptation to changing environments is a continuous process with wide-ranging consequences. Our study reveals that laboratory domestication has profoundly shaped the phenotypic and transcriptional landscape of *C. elegans*. While the N2 Lab strain showed moderate and pre-conditioned responses to several environmental challenges, the wild strains (the N2 Ancestral strain and a recent tropical isolate from Mexico City), exhibited more dynamic and stress-specific transcriptional responses, reflecting their divergent evolutionary trajectories.

Phenotypically, the Mexican strain displayed reduced lifespan and fertility, but enhanced resistance to *P. aeruginosa*, while the Ancestral strain showed high resistance to oxidative stress and low thermotolerance. These life-history traits correlate with transcriptional signatures, including upregulation of immune-related genes in the Mexican strain and oxidative stress genes in the Ancestral strain. Such divergence reflects life-history trade-offs likely shaped by natural versus artificial environments; for example enhanced pathogen resistance in the Mexican strain may be compromising its resistance to oxidative stress.

Remarkably, we found that the Ancestral and Mexican strains share greater transcriptional similarity with each other than with the N2 Lab strain, despite the Ancestral shared origin with the latter. This suggests that laboratory domestication, over thousands of generations, has led to a widespread transcriptional divergence alongside reduced plasticity in the N2 lab strain. This is exemplified by the elevated expression of stress-related genes in the N2 Lab even at 20 °C, possibly reflecting a constitutive stress adaptation to laboratory conditions. In this sense, we could argue that laboratory conditions are in fact stressful for the wild isolates: for instance, exposure to to atmospheric oxygen concentration or changes in diet, temperature variability and pathogen exposure, that have demonstrated to lead to wide changes in gene expression in *C. elegans* (Gómez-Orte et al. 2017; Jovic et al. 2019; Lansdon et al. 2022; Branicky et al. 2022).

At the transcriptional level, differential expression analysis highlighted cuticle and immune related genes as prominent axes of divergence. The wild strains upregulated numerous collagen genes, likely reflecting adaptive remodeling of the cuticle in response to environmental challenges. Furthermore, downregulation of chromatin remodelers in Mexican strain alongside with upregulation of histone acetylation reader orthologs, suggest that epigenetic mechanisms may contribute to strain-specific transcriptional response.

In response to acute heat stress, all strains showed distinct transcriptional responses albeit activation of a conserved core of the HSR program (i.e. *hsp-16.2, hsp-16.41, hsp-70*). The Mexican and Ancestral strain shared additional responses involving *hsf-1*, *daf-2*, *daf-18*, and *skn-1*, which were unresponsive in the N2 Lab. These results suggest a more flexible inducible heat response in wild strains (at 2-hour exposure), in contrast to pre-adapted, perhaps a less dynamic response in the N2 Lab. The Ancestral-specific heat induction of mitochondrial UPR components (hsp-6, hsp-60) and differential regulation of ROS tolerance genes (*pah-1*, *sod-3*, *ctl-1*) further illustrate distinct strategies employed by each lineage.

Extensive genetic and phenotypic comparisons between the N2 Lab strain and other wild isolate strains have been widely performed highlighting the N2 Lab diversification (Volkers et al. 2013; Viñuela et al. 2012; Kamkina et al., 2016; Snoek et al. 2017; Snoek et al. 2019; Snoek et al. 2021), however, to our knowledge, this is the first study to also include both the N2 Ancestral strain as well as a recently isolated wild strain. And, although genetic variations may be also contributing to the observed differences, other layers of regulation, such as epigenetic differences, would have to be explored as well.

Altogether, our findings underscore the importance of considering strain history and domestication in experimental designs. The inclusion of both, an ancestral and a recently wild strain, allowed us to bridge the gap between long-term laboratory adaptation and natural variation. Future work should explore stress response dynamics over time and assess epigenetic profiling, including histone modifications and chromatin state mapping, that may underlie regulatory mechanisms contributing to the observed differences between these three evaluated strains. By continuing to integrate wild isolates into functional studies, we can gain a more complete understanding of how *C. elegans* adapts to diverse environments at molecular, physiological, genetic and epigenetic levels.

## Materials and Methods

### Nematode and bacterial strains

*C. elegans* strains used in this study were: wild-type N2 (var. Bristol) and the N2 Ancestral strain, originally frozen in 1969 and thawed in 1980, estimated to be fewer than six generations from the original stock; both strains were obtained from the Caenorhabditis Genetics Center (CGC). The VJV001 (Mexican) strain was isolated in Mexico City in June 2023 (this study). Strains were maintained on NGM plates, fed with *E. coli* OP50-1 at 20 °C (Stiernagle 2006) and synchronized by bleaching. The following bacterial strains were used: *Pseudomonas aeruginosa* strain PA14; and *E. coli* RNAi clones targeting *dpy-11* and *pos-1* from the Ahringer RNAi library (Kamath and Ahringer 2003). *C. elegans* protocols were approved by the IFC-IACUC (protocol VVR191-22)

### Mexican *C. elegans* isolation

Sampling was conducted in the gardens of Instituto de Fisiología Celular, UNAM, in México City (coordinates: 19.32840, -99.17846). Pieces of ripe pear were buried 1-2 cm deep in moist soil and collected after one week along with surrounding soil and debris. Wild isolates were processed as specified in Barrière and Félix (2014). Briefly, each sample was transferred to a 100 mm NGM plate seed with *E. coli* lawn and sealed with parafilm. After 3-5 days plates were screened for worm’s trails; adult worms were individually transferred to a 30 mm plate with food. After 3 days, the presence of double-bulb pharynx, characteristics of *C. elegans* was confirmed by microscopy. For individuals showing this feature, species identity was confirmed by PCR amplicon of the ITS2 gene (Internally Transcribed Spacer between 5.8S and 28S rDNA genes) using RHAB1350F (5’-TACAATGGAAGGCAGCAGGC) and RHAB1868R (5’-CCTCTGACTTTCGTTCTTGATTAA) primers that in *C. elegans* gives an 557 bp amplicon. Samples yielding this product were subjected to a second PCR with primers 5.8S-1 (5’-CTGCGTTACTTACCACGAATTGCARAC) and KK-28S-22 (5’CACTTTCAAGCAACCCGAC), which results in a 777 bp amplicon specific for *C. elegans*. The amplicon was sequenced by Sanger and analyzed by NCBI/BLAST tool.

### Lifespan assay

Assays were conducted using synchronized populations of L4-stage worms. For each strain, approximately 180 worms were distributed in three 30 mm plates and monitored daily for survival until all individuals had died. During the first 5–7 days, worms were transferred to fresh plates daily. After this period, worms were transferred every other day to ensure food supply. Worms that exhibited bagging or crawled off the plate were censored from the analysis. Worms were considered dead if they failed to respond to gentle prodding with a platinum wire. Each lifespan assay was repeated in three independent biological replicates.

### *Pseudomonas aeruginosa* PA14 survival assay

A fast-killing *Pseudomonas aeruginosa* PA14 assay was performed as described by Liu et al. (2024), with minor modifications. Briefly, PA14 cultures were initiated from fresh LB plates (less than one month). Prior to the assay, bacteria were grown overnight in LB broth for no longer than 16 hours. 150 μL of the overnight culture was spread evenly onto 33 mm NGM plates, ensuring complete surface coverage. Plates were allowed to dry and incubated upside down at 37°C for 15 hours followed by 6 hours incubation at 26°C. Approximately 50 synchronized L4-stage worms were transferred to each plate (3 plates per strain) and maintained at 26°C. Worm mortality was assessed at intervals of 8-15 hours over ∼50 hours. The assay was repeated in twice independent biological replicates.

### Fertility assays

Synchronized L4-stage worms were individually placed on 33 mm NGM plates seeded with *E. coli*. Worms were transferred to fresh plates every 24 hours. for six days. Progeny were counted for 48 hours. after removal of parental hermaphrodite. This procedure was repeated until the progeny of ≥ 18 parental worms were analyzed. Total fertility was calculated by summing the number of hatched embryos (L1 larvae) produced by each worm.

### 35 °C survival assay

One-day adult worms (approximately 25 per plate) were placed on 33 mm NGM plates seeded with *E. coli*. The experiment was conducted in triplicate, with three plates per strain. Plates were incubated in a water bath at 35°C, one plate per strain was removed each hour from 4 to 9 hours. After removal, plates were transferred to 20°C for recovery. Survival was assessed 12 hours after the final time point by scoring live and dead worms.

### Oxidative stress resistance assay

Synchronized L4-stage worms were exposed to oxidative stress using paraquat. Approximately 30 individuals per strain were placed into each well of a 96-well plate containing 300 μL of M9 buffer supplemented with *E. coli* and 250 mM of paraquat. Each strain was tested in duplicate. Worm mortality was recorded every 30 minutes over the course of the experiment.

### Chemotaxis

Synchronization was performed as described by Remy and Hobert (2005): day 1 adults were allowed to lay approximately 300 eggs in 60 mm NGM plates with food. Adults were discarded by gently washing the plates with M9; remaining eggs were incubated at 20 °C until they reached the young adult stage. Chemotaxis assays were conducted as in Bargmann et al. (1993): approximately 200 worms were placed in the center of a 100 mm plate containing the test odorants at one end of the plate and the vehicle, as a control, at the other. NaN_3_ was placed on both ends to paralyze the animals. The odorant concentrations and vehicles used were the following: benzaldehyde 0.33% diluted in H20; butanone 0.01% diluted in 100% ethanol; isoamyl alcohol 1% diluted in 100% ethanol; and octanol 0.1% diluted in 100% ethanol. Chemotaxis index was calculated with the following formula: *CI* = (*N odorant* − *N control*) / *N total*

### RNA interference

RNAi experiments targeting *pos-1* and *dpy-11* genes were conducted using *E. coli* HT115 strain engineered to express double-stranded RNA (Kamath and Ahringer 2003). Bacteria were cultured overnight at 37 °C in LB broth containing ampicillin (100 mg/ml) and tetracycline (12.5 mg/ml). NGM plates with ampicillin (10 ug/mL) were overlaid with IPTG at desired concentration to induce dsRNA expression. Plates were left overnight at RT° protected from light. For *pos-1* RNAi, one L4-larva was placed on each RNAi plate and progeny were monitored daily for six days. To assess gene silencing, hatched and unhatched progeny was counted. For *dpy-11*,one gravid worm was placed per plate for ∼4 hours. to lay eggs and synchronize the population. Scoring was counted 72 hours later to asees dumpy phenotype. All RNAi experiments were performed at 20 °C

### RNA-seq library preparation

Total RNA was isolated by collecting worms with M9 buffer and washed twice to remove bacteria. 300 μL of Trizol were added to each sample followed by 8 cycles of LN2 freezing and thawing at 37 °C to ensure lysis. This was followed by 4 cycles of vortexing (20 sec. at max speed, then 20 sec set in ice). Samples were centrifuged at 10,000 xg for 1 min. Then RNA was purified using Direct-zol RNA MicroPrep (Zymo, R2062) following manufacturer’s instructions. RNA’s integrity and concentration were evaluated on 2100 Bioanalyzer (Agilent). Only samples with a RIN >8 were selected for library preparation. Libraries were prepared from 700 ng of RNA using the Nextera XT DNA library prep kit (Illumina) following the reference guide. Synthesized libraries were sequenced on NextSeq 500 System (Illumina) using paired-end 75-bp reads.

### RNA-seq data analysis

Data quality was assessed using FastQC. Alignments of paired-end samples were generated using STAR (version=2.7.11b) with default parameters, except for “--sjdbOverhang 149” and “--genomeSAindexNbases 12”, using the *C. elegans* genome WBcel235 from the Ensembl portal. Raw count matrices were generated using FeatureCounts with flags “-p -a”. Differential gene expression was analyzed with DESeq2 (v1.42.1). Genes with an adjusted *P value* < 0.05 and log_2_ fold change > 2 and < -2 were considered differentially expressed, unless specified otherwise. Pathway enrichment was performed using the WormCat web tool (Holdorf et al. 2020) with a *P*-value cutoff of 0.05 to define enriched categories. Heatmaps were generated using DESeq2 variance-stabilizing transformation normalized counts and the R package Pheatmap. Volcano plots and transcript expression patterns were generated using RStudio (v4.3.3) using custom code available at [https://github.com/jaq77/Mexa-C.-elegans.git]. Genes lists for heatmaps were obtained from amigo.geneontology.org/amigo, (Ashburner et al. 2000; Gene Ontology Consortium 2023) using the Advanced annotation search, with the following fillers: Organism: Caenorhabditis elegans; Type: gene; GO class (including “regulates”). GO class for: (1) Lifespan = Insulin, Energy metabolism, Determination of adult lifespan; (2) Fertility = Spermatogenesis, Spermatid development, Spermidine biosynthetic process, Amoeboid sperm motility, Flagellated sperm motility, Sperm flagellum, Positive regulation of amoeboid sperm motility, Sperm ejaculation, Sperm entry, Spermathecum morphogenesis, Spermidine metabolic process, Gonad development, Gonad morphogenesis, Positive regulation of gonad development, Gonadal mesoderm development, Male gonad development, Female gonad development, Positive regulation of female gonad development, Oogenesis, Positive regulation of oogenesis, Oocyte maturation, Oocyte development, Positive regulation of oocyte development, Oocyte growth, Regulation of meiotic cell cycle process involved in oocyte maturation, Oocyte morphogenesis, Positive regulation of oocyte maturation, Negative regulation of oocyte maturation, Regulation of oocyte development; (3) ROS = Response to oxidative stress, Positive regulation of response to oxidative stress, Cellular response to oxidative stress, Negative regulation of response to oxidative stress, Regulation of response to oxidative stress, Cell redox homeostasis, some genes from Rafikova et al. (2020); (4) Heat stress = Response to heat, Heat acclimation, Heat shock protein binding, Cellular response to heat, Negative regulation of cellular response to heat, Positive regulation of cellular response to heat, Regulation of cellular response to heat, Cellular heat acclimation; (5) Immune system = Immune response, Antibacterial innate immune response, Defense response to gram positive bacterium, Defense response to gram negative bacterium; (6) Locomotion = Locomotion, Muscle contraction, Response to stimulus; (7) Nervous system = Nervous system development; (8) Epigenetic = Epigenetic regulation, Chromatin remodeling, Chromosome organization, Histone binding, Histone acetyltransferase activity, Histone deacetylase activity, Methylated histone binding.

## Data Availability

*The data underlying this article are available in* Gene Expression Omnibus repository at [URL], *and can be accessed with* [unique identifier, e.g. accession number, deposition number]. NOTE: the data will be released upon acceptance of the manuscript or by request of the reviewers.

## Acknowledgements

J Hersch-González conducted this study to fulfilll the requirements of the Programa de Doctorado en Ciencias Bioquímicas of the Universidad Nacional Autónoma de Meéxico (UNAM), and received a doctoral scholarship from the Consejo Nacional de Humanidades, Ciencias y Tecnologías, CONACYT (now Secretaría de Ciencia, Humanidades, Tecnología e Innovación, SECIHTI) (#CVU 846476). J Hersch-González was supported by a CONAHCYT (now SECIHTI) scholarship (788519). This project was supported by the PAPIIT-UNAM grant IN217824 to VJV. At IFC, we thank the UBM: Laura Ongay-Larios, Guadalupe Codiz and Minerva Mora the UBMI; Augusto César Poot-Hernándezt and Carlos Peralta, and U de Computo, Imagenología and Biblioteca. We gratefully acknowledge the Caenorhabditis Genetics Center (CGC) for providing the strains used in this work. The CGC is funded by the NIH Office of Research Infrastructure Programs (P40 OD010440). We thank Rosa Navarro and Diego Cortez for their valuable contributions throughout the conduct of this study.

