## Supplemental files for "Phenotypic and transcriptomic similarity between the N2 Ancestral and a tropical wild isolate of *C. elegans* reveals divergence from the reference Bristol strain"

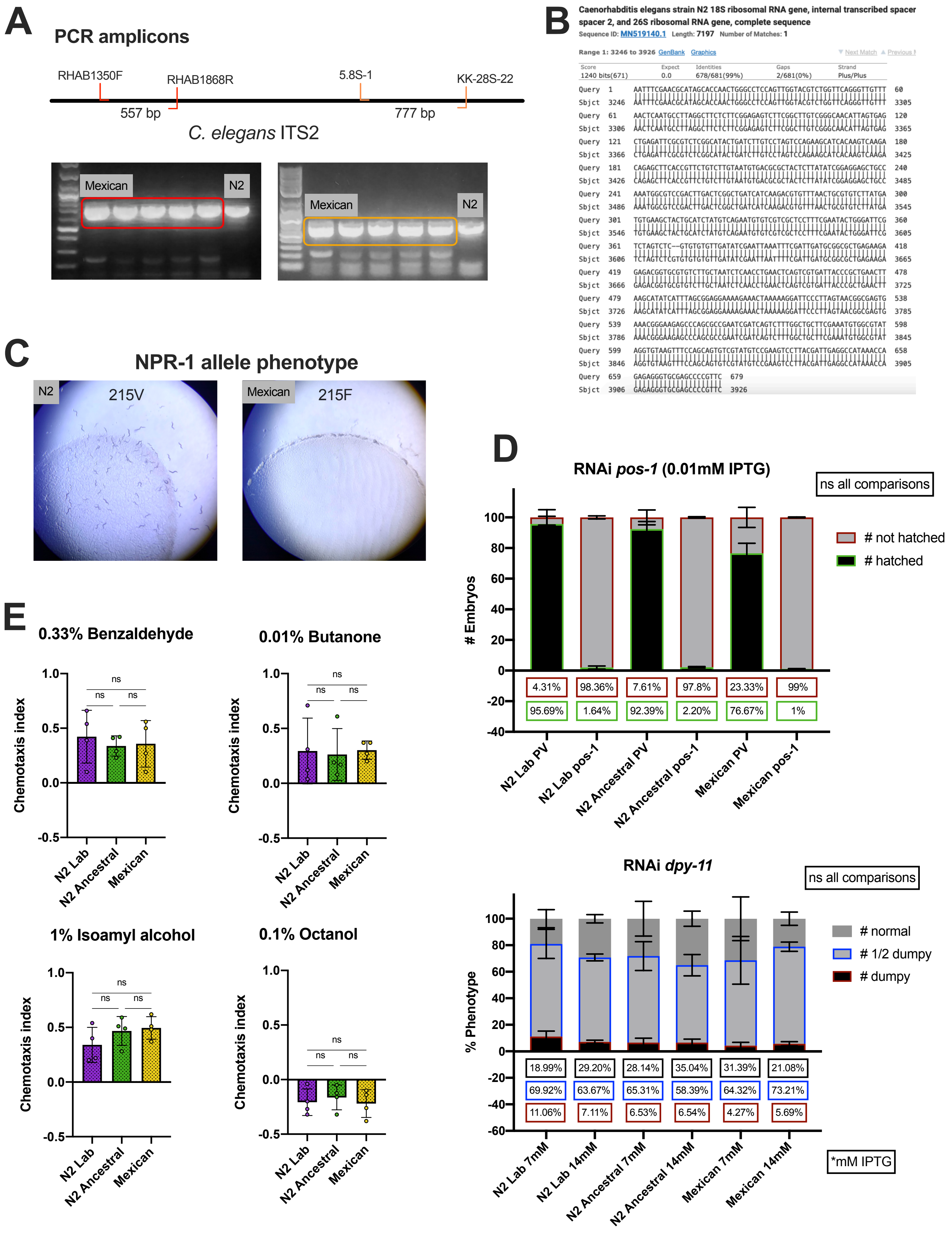

A
PCR amplicons
RHAB1350F
RHAB1868R
5.8S-1
KK-28S-22
557 bp
777 bp
C. elegans ITS2
Mexican
N2
Mexican
N2
B
Caenorhabditis elegans strain N2 18S ribosomal RNA gene, internal transcribed spacer 2, and 26S ribosomal RNA gene, complete sequence
Sequence ID: MN519140.1 Length: 7197 Number of Matches: 1
Range 1: 3246 to 3926 GenBank Graphics
Score 1240 bits(671)
Expect 0.0
Identities 678/681(99%)
Gaps 2/681(0%)
Strand Plus/Plus
C
NPR-1 allele phenotype
N2
215V
Mexican
215F
D
RNAi pos-1 (0.01mM IPTG)
ns all comparisons
### Embryos
### not hatched
### hatched
4.31%
98.36%
7.61%
97.8%
23.33%
99%
95.69%
1.64%
92.39%
2.20%
76.67%
1%
N2 Lab PV
N2 Lab pos-1
N2 Ancestral PV
N2 Ancestral pos-1
Mexican PV
Mexican pos-1
RNAi dpy-11
ns all comparisons
% Phenotype
### normal
### 1/2 dumpy
### dumpy
18.99%
29.20%
28.14%
35.04%
31.39%
21.08%
69.92%
63.67%
65.31%
58.39%
64.32%
73.21%
11.06%
7.11%
6.53%
6.54%
4.27%
5.69%
N2 Lab 7mM
N2 Lab 14mM
N2 Ancestral 7mM
N2 Ancestral 14mM
Mexican 7mM
Mexican 14mM
*mM IPTG
E
0.33% Benzaldehyde
0.01% Butanone
1% Isoamyl alcohol
0.1% Octanol
Chemotaxis index
ns
N2 Lab
N2 Ancestral
Mexican


Figure S1.

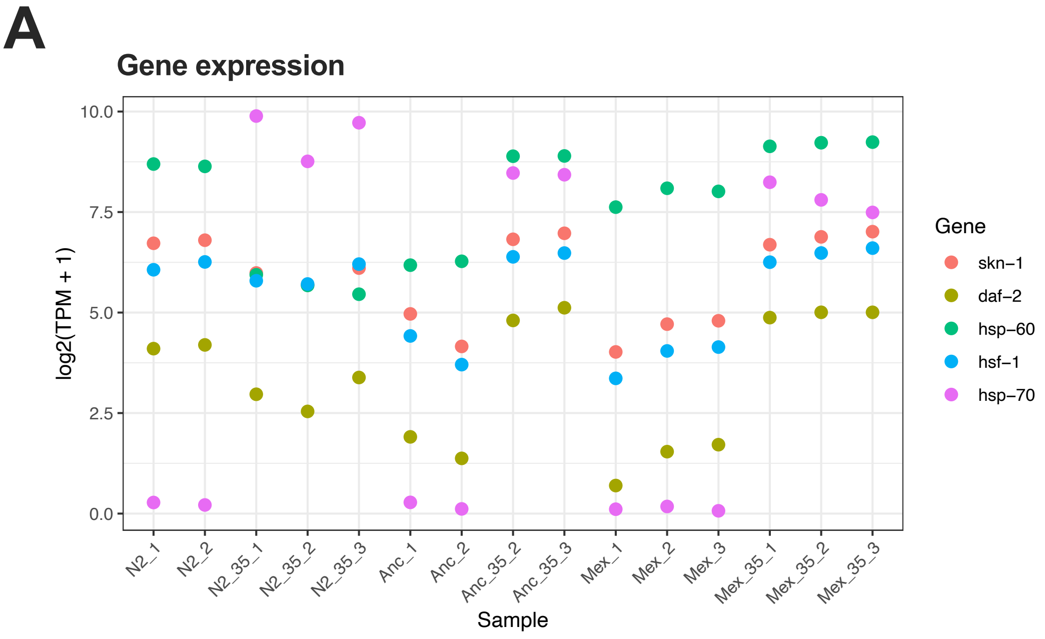
A
Gene expression
10.0
7.5
5.0
2.5
0.0
log2(TPM + 1)
N2_1
N2_2
N2_35_1
N2_35_2
N2_35_3
Anc_1
Anc_2
Anc_35_2
Anc_35_3
Mex_1
Mex_2
Mex_3
Mex_35_1
Mex_35_2
Mex_35_3
Sample
Gene
skn−1
daf−2
hsp−60
hsf−1
hsp−70

**Figure S2.**

**Table S1** Enriched Gene Ontology Terms for 20 °C expressed transcripts

| | Number of transcripts | Gene Ontology Term[a] | Number of genes | Adjusted *P*-value (Fisher-Bonferroni)[b] |
|---|---|---|---|---|
| **N2 Lab & N2 Ancestral** | 209 | Unassigned: BTB and MATH domain | 9 | $9.12x10^{-11}$ |
| | | Proteolysis proteasome: E3: F box | 16 | $1.06x10^{-5}$ |
| | | Unassigned: regulated by multiple stresses | 32 | $8.46x10^{-5}$ |
| | | Proteolysis proteasome: ubiquitin | 3 | $3.20x10^{-3}$ |
| **N2 ancestral only** | 46 | Proteolysis proteasome: E3: F box | 17 | $1.67x10^{-16}$ |
| | | Transmembrane transport: potassium channel | 4 | $4.69x10^{-4}$ |
| | | Stress response: detoxification: ugt | 3 | $1.95x10^{-3}$ |
| **N2 Ancestral & Mexican** | 691 | Proteolysis proteasome: E3: F box | 83 | $1.07x10^{-41}$ |
| | | Unassigned: regulated by multiple stresses | 97 | $1.49x10^{-12}$ |
| | | Metabolism: insulin | 14 | $6.09x10^{-10}$ |
| | | Transcription factor: homeodomain | 16 | $4.03x10^{-7}$ |
| | | Transcription factor: T box | 8 | $1.47x10^{-5}$ |
| | | Extracellular material: cuticlin | 8 | $3.78x10^{-4}$ |
| | | Neuronal function: synaptic function: neuropeptide | 14 | $7.81x10^{-4}$ |
| | | Transmembrane protein: unassigned | 68 | $1.08x10^{-3}$ |
| | | Extracellular material: collagen | 15 | $3.80x10^{-3}$ |
| **Mexican only** | 43 | Proteolysis proteasome: E3: F box | 6 | $1.11x10^{-3}$ |

[a] Category 3 from WormCat
[b] FDR-adjusted *P-value* is a measure of enrichment (Fisher exact test; Bonferroni Correction) of the GO term

**Table S2** Differential gene expression $log_2$ fold change (LFC) values

| Gene name | 20 ºC | | | 35 ºC | | |
|---|---|---|---|---|---|---|
| | N2 Ancestral vs. N2 Lab | Mexican vs. N2 Lab | Mexican vs. N2 Ancestral | N2 Lab | N2 Ancestral | Mexican |
| *C04G2.10* | - | -1.78 | -2.42 | - | - | - |
| *C25F9.4* | -2.10 | -8.52 | -6.42 | -4.97 | 1.87 | - |
| *C25F9.5* | -2.82 | -8.67 | -5.84 | -6.42 | 2.40 | - |
| *cbp-2* | - | -9.91 | -8.85 | -3.93 | - | - |
| *cct-2* | -1.66 | - | - | - | - | - |
| *cct-4* | -1.56 | - | - | - | - | - |
| *cct-8* | -1.88 | - | - | - | 1.73 | - |
| *ctl-1* | - | -8.36 | -7.99 | -1.62 | - | - |
| *daf-16* | -1.65 | -1.46 | - | - | 1.92 | 1.70 |
| *daf-18* | -2.49 | -2.84 | - | - | 2.49 | 3.27 |
| *daf-2* | -2.65 | -3.04 | - | - | 3.38 | 3.90 |
| *dnj-25* | -1.59 | - | - | - | 2.00 | 1.86 |
| *dnj-27* | 1.51 | - | - | 2.95 | -1.63 | - |
| *dnj-5* | -1.94 | -2.40 | - | - | 2.01 | 2.69 |
| *elt-3* | 1.66 | 1.14 | - | - | - | - |
| *endu-2* | -2.02 | - | 1.60 | -3.41 | 2.19 | - |
| *F08H9.3* | - | - | - | 8.45 | - | - |
| *F44E5.5* | - | - | - | 5.63 | 5.84 | - |
| *gpx-5* | 1.94 | - | - | - | -1.95 | -1.60 |
| *gst-10* | 1.72 | - | - | -1.80 | -3.46 | -1.94 |
| *gst-4* | 1.75 | - | - | - | -2.64 | -2.15 |
| *hil-2* | 2.28 | 2.01 | - | 2.23 | -2.53 | -1.80 |
| *hsf-1* | -1.83 | -2.01 | - | - | 1.97 | 2.32 |
| *hsp-1* | -2.11 | - | - | -2.32 | 1.97 | - |
| *hsp-110* | -2.56 | -1.53 | - | - | 3.63 | 2.31 |
| *hsp-12.2* | 3.08 | 3.26 | - | 2.25 | -2.20 | -3.47 |
| *hsp-12.3* | - | 3.47 | 3.03 | - | - | -1.71 |
| *hsp-12.6* | - | - | - | 2.38 | 3.61 | - |
| *hsp-16.2* | - | - | - | 14.21 | 9.75 | 9.27 |
| *hsp-16.41* | - | - | - | 17.56 | 8.79 | 8.01 |
| *hsp-6* | -2.51 | - | - | -2.00 | 2.72 | 1.50 |
| *hsp-60* | -2.10 | - | 1.70 | -2.74 | 2.20 | - |
| *hsp-70* | - | - | - | 12.19 | 10.88 | 11.05 |
| *hsp-90* | -1.62 | - | - | - | 1.58 | - |
| *jmjc-1* | - | - | - | 3.06 | -2.64 | -1.62 |
| *jnk-1* | - | - | - | - | - | -2.10 |
| *K07F5.16* | - | - | - | 2.78 | - | - |
| *lipl-4* | - | 1.86 | 2.63 | - | - | - |
| *lys-7* | - | 3.09 | 2.24 | - | - | -2.06 |
| *M04C3.1* | -2.60 | -10.54 | -7.94 | -3.50 | 2.95 | - |
| *M04C3.2* | -2.62 | -7.49 | -4.87 | -7.11 | 2.76 | - |
| *mtl-2* | - | - | - | -3.91 | -3.40 | -2.61 |

| | | | | | | |
|---|---|---|---|---|---|---|
| *pah-1* | 2.14 | - | - | - | -2.12 | -1.58 |
| *rme-8* | -2.65 | -2.18 | - | -1.90 | 2.93 | 2.47 |
| *rpn-2* | -1.84 | - | - | - | 1.78 | - |
| *skn-1* | -1.90 | -1.92 | - | - | 1.90 | 2.02 |
| *sod-3* | 2.68 | - | -2.26 | - | -2.09 | - |
| *spl-2* | - | -2.58 | -2.67 | -4.53 | -2.86 | - |
| *swan-1* | - | - | - | 2.82 | - | - |
| *ttx-1* | 2.07 | 2.48 | - | - | -2.41 | -2.89 |
| *uba-1* | -2.48 | -1.74 | - | - | 2.44 | 1.97 |
| *Y17G9B.8* | -2.33 | - | 3.32 | - | 2.28 | - |

**Table S3** Enriched Gene Ontology Terms for 35 °C expressed transcripts

| | Number of transcripts | Gene Ontology Term[a] | Number of genes | Adjusted *P*-value (Fisher-Bonferroni)[b] |
|---|---|---|---|---|
| **N2 Lab only** | 390 | Proteolysis proteasome: E3: F box | 84 | $6.10x10^{-60}$ |
| | | Transcription factor: homeodomain | 12 | $9.56x10^{-7}$ |
| | | Transcription factor: forkhead | 4 | $4.14x10^{-3}$ |
| | | Transcription factor: bHLH | 5 | $9.99x10^{-3}$ |
| **N2 Lab & N2 Ancestral** | 460 | Unassigned: regulated by multiple stresses | 104 | $1.05x10^{-26}$ |
| | | Cytoskeleton: microtubule: tau tubulin kinase | 14 | $4.73x10^{-9}$ |
| | | Major sperm protein | 10 | $3.17x10^{-8}$ |
| | | Unassigned: BTB and MATH domain | 7 | $3.31x10^{-5}$ |
| **N2 Ancestral only** | 490 | Unassigned: regulated by multiple stresses | 150 | $1.34x10^{-51}$ |
| | | Signaling: phosphatase: Y | 38 | $3.91x10^{-35}$ |
| | | Cytoskeleton: microtubule: tau tubulin kinase | 26 | $2.05x10^{-22}$ |
| | | Transmembrane protein: unassigned | 85 | $5.34x10^{-16}$ |
| | | Signaling: Y kinase | 15 | $5.73x10^{-10}$ |
| | | Signaling: phosphatase: unassigned | 11 | $8.14x10^{-5}$ |
| **N2 Ancestral & Mexican** | 309 | Stress response: C-type Lectin | 16 | $2.02x10^{-6}$ |
| **Mexican only** | 31 | Proteolysis general: lysozyme | 2 | $1.40x10^{-3}$ |
| **N2 Lab & Mexican** | 184 | Proteolysis proteasome: E3: F box | 47 | $3.84x10^{-37}$ |
| | | Transcription factor: T box | 5 | $2.41x10^{-5}$ |

[a] Category 3 from WormCat

[b] FDR-adjusted *P-value* is a measure of enrichment (Fisher exact test; Bonferroni Correction) of the GO term

**Figure S1. Species confirmation, RNA interference response and and chemotaxis assays in the Mexican strain**

**(A)** Map of the ITS2 gene (Internally Transcribed Spacer between 5.8S and 28S rDNA genes) of *C. elegans* and PCR amplification of ITS2 fragments: 557 bp using RHAB1350F and RHAB1868R primers; 777 bp with 5.8S-1 and KK-28S-22 primers. **(B)** Alignment of the Sanger-sequenced 777 bp amplicon using the NCBI/BLAST tool. Sequence aligns 100% with the *C. elegans* reference genome **(C)** Population feeding behavior: Left, the N2 Lab strain shows the *npr-1* 215V 'solo' feeding phenotype; Right, the Mexican strain shows the *npr-1* 215F 'social' feeding phenotype. **(D)** RNA interference (*RNAi*) assay. Embryonic lethality was assessed by silencing the *pos-1* gene with 0.01mM IPTG. The *Dumpy* phenotype was induced by silencing the *spy-11* gene using 7 mM and 14 mM IPTG. No differences were observed between the three strains in either assay. Statistically significance was assessed by two-way ANOVA with multiple comparisons. Graphs show representative experiments with consistent trends across 3-4 independent replicates. **(E)** Chemotaxis assays towards benzaldehyde (0.33%), butanone (0.01%), isoamyl alcohol (1%) and octanol (0.1%) showed no significant differences among the three strains. Statistical significance was assessed by one-way ANOVA with multiple comparisons.

**Figure S2. *skn-1*, *daf-2*, *hsp-60*, *hsf-1* and *hsp-70* gene expression at 20 ºC and 35 ºC.**
**(A)** Transcript per million (TPM) and shown per sample per strain: N2 = N2 Lab 20 ºC; N2_35 = N2 Lab 35 ºC; Anc = N2 Ancestral 20 ºC; Anc_35 = N2 Ancestral 35 ºC; Mex = Mexican 20 ºC; Mex_35 = Mexican 35 ºC.
